# CCSer2 gates dynein activity at the cell periphery

**DOI:** 10.1101/2024.06.13.598865

**Authors:** Juliana L. Zang, Daytan Gibson, Ann-Marie Zheng, Wanjing Shi, John P. Gillies, Chris Stein, Catherine M. Drerup, Morgan E. DeSantis

## Abstract

Cytoplasmic dynein-1 (dynein) is a microtubule-associated, minus end-directed motor that traffics hundreds of different cargos. Dynein must discriminate between cargos and traffic them at the appropriate time from the correct cellular region. How dynein’s trafficking activity is regulated in time or cellular space remains poorly understood. Here, we identify CCSer2 as the first known protein to gate dynein activity in the spatial dimension. CCSer2 promotes the migration of developing zebrafish primordium cells and of cultured human cells by facilitating the trafficking of cargos that are acted on by cortically localized dynein. CCSer2 inhibits the interaction between dynein and its regulator Ndel1 exclusively at the cell periphery, resulting in localized dynein activation. Our findings suggest that the spatial specificity of dynein is achieved by the localization of proteins that disinhibit Ndel1. We propose that CCSer2 defines a broader class of proteins that activate dynein in distinct microenvironments via Ndel1 inhibition.

## Introduction

Microtubules, microtubule-associated proteins, and microtubule motors form a complex system of proteins that extend throughout the cytoplasm of most eukaryotic cells, facilitating molecular exchange between distinct cellular regions.^1^ The activities of motor proteins must be dynamically controlled for the system to support intracellular trafficking. For example, to move cargo, motors must receive and integrate signals that relay identity, spatial, and temporal information. In other words, motor machinery must decipher if the right cargo is being moved from the correct region of the cell at the appropriate time.

There are two classes of microtubule-associated motors: kinesins and cytoplasmic dynein-1 (dynein, hereafter).^2,3^ To properly traffic cargo, both classes of motors must robustly and accurately respond to spatiotemporal cues. There are over twenty different kinesins that traffic cargo during interphase.^2,4^ These kinesins have diverged and specialized to meet the trafficking requirements of different kinds of cargo, which means that kinesins’ ability to interpret spatiotemporal information about cargo is, in part, determined at the protein sequence level.^2^ In contrast, dynein– as the primary minus-end directed microtubule motor– traffics every cargo that moves in the retrograde direction.^3,5^ Therefore, dynein’s ability to traffic cargo with specificity is not encoded within dynein’s protein sequence and is instead conferred by interaction with dynein regulators.

We have a good understanding of how cargo-identity information is conveyed to dynein. To transport cargo, dynein must bind to the multi-subunit complex, dynactin, as well as a protein called an activating adaptor (adaptor, hereafter).^5^ Together, dynein-dynactin-adaptor complexes form an active transport complex that can move processively along microtubules (Figure S1A).^6,7^ There are *∼*20 confirmed or putative adaptors, which associate with distinct cargo types and thus convey cargo-identity information to dynein.^5^ However, many adaptors localize in close or overlapping cellular regions and therefore do not convey spatial information to the dynein motor.^5^ Finally, we know very little about how dynamic cellular events influence dynein’s association with cargo, so it is unclear how dynein activity is regulated temporally. As such, our understanding of dynein regulation is one-dimensional; we lack information about how dynein activity is gated in both space and time.

Together, dynein, dynactin, and adaptors represent the core dynein trafficking machinery. The proteins Lis1 and Ndel1 (and its paralogue, Nde1) constitute a second sphere of regulatory factors that have two key functions (Figure S1A).^8,9^ First, Lis1 and Ndel1 support dynein localization to many cellular regions, including the microtubule plus-end, the nuclear envelope, and the kinetochore.^10–16^ Importantly, Lis1 and Ndel1 promote dynein localization to multiple cellular microenvironments in a way that is likely independent of adaptor or cargo localization. Second, Lis1 and Ndel1 modulate dynein activation by controlling dynein’s affinity for dynactin: Lis1 favors dynein activation by driving dynein-dynactin-adaptor binding, while Ndel1 inhibits dynein activation by binding competitively with dynactin and by sequestering Lis1 away from dynein’s motor.^17–23^ Lis1 and Ndel1 bind with high affinity and, despite having opposing effects on dynein activation separately, likely function in tandem to regulate dynein motility in cells (Figure S1A).^20,21,23,24^

To understand dynein function, we must identify additional factors that regulate dynein’s ability to traffic cargo with spatial or temporal specificity. Because Lis1 and Ndel1 modulate both dynein activity and localization in cellular space, we hypothesized that Lis1 and Ndel1 are likely nodes through which spatial information is transmitted to dynein and set out to identify and characterize the activity of Lis1 or Ndel1 binding partners. To do this, we used proximity-dependent biotinylation coupled to mass-spectrometry (MS) to establish the interactome of Lis1 and Ndel1. We identified a poorly characterized protein, CCSer2, which we find binds to Ndel1 directly. We find that depletion of CCSer2 causes dramatic migration defects, both in developing zebrafish embryos and in cell culture. Unexpectedly, we find that CCSer2 function is not broadly acting, but instead appears to promote dynein activity only at the cell periphery. The outcome of this is that CCSer2 depletion causes aberrant dynein-driven centrosome positioning and early endosomes trafficking, which are both cargos that are operated on by cortically localized dynein. We also establish that CCSer2 functions at the molecular level to promote dynein motility by attenuating Ndel1-mediated inhibition of dynein activation. Our work shows that CCSer2 functions to inhibit Ndel1-mediated inhibition of dynein exclusively at the cell edge to locally activate populations of dynein. CCSer2 activity represents a novel regulatory gate that conveys information about cellular location, rather than cargo identity, to the dynein motor. These findings deepen our understanding of how dynein activity can be deployed locally to promote cargo trafficking specificity.

## Results

### CCSer2 interacts with Lis1 and Ndel1 by binding directly to Ndel1

To identify proteins that modulate dynein’s response to spatial cellular cues, we determined the interactome of Lis1 and Ndel1 using proximity-dependent biotinylation coupled to mass spectrometry (MS).^25,26^ We fused the promiscuous biotin-ligase tag, BioID, to Lis1 or Ndel1 and generated stable HEK-293 cell lines. For these experiments, we appended BioID to the β-propellor at the C-terminus of Lis1 as this is the domain that Lis1 uses to interact with most of its known binding partners. Because Ndel1 is a long coiled coil and interacts with proteins using domains that are found throughout the protein, we generated two Ndel1 constructs: one with BioID on the N-terminus and one with BioID on the C-terminus.^27^ After culturing cells in the presence of biotin for 16 hours, we performed a streptavidin immunoprecipitation followed by MS to identify near neighbors of both proteins. Many known and well-characterized binding partners of Lis1 and Ndel1 were enriched at levels above the BioID control in each dataset, which gave us confidence that our screen could identify bona fide interactors of Lis1 and Ndel1 (Figure 1A; S1B-D).

**Figure 1.**
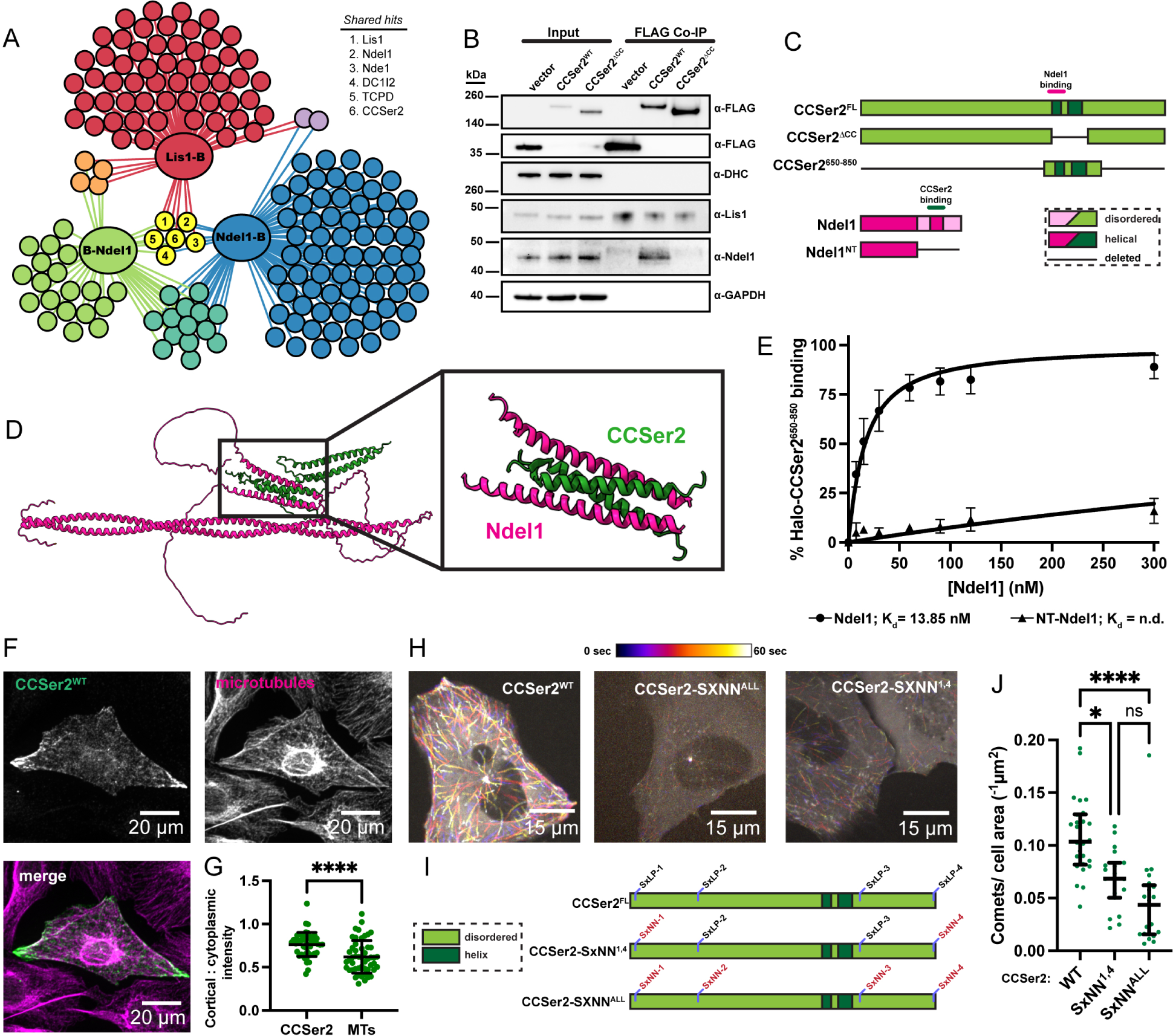
CCSer2 is in the interactome of Lis1 and Ndel1 via a direct binding interaction with Ndel1’s C-terminal coiled coil. A) Significant and enriched hits shown as spheres and color-coded according to the data set they were found in: Lis1-BioID (red), Ndel1-BioID (green), BioID-Ndel1 (blue), Lis1-BioID/BioID-Ndel1 (orange), Lis1-BioID/Ndel1-BioID (purple), BioID-Ndel1/Ndel1-BioID (teal), or all data sets (yellow). The identity of the hits found in all datasets is indicated and discussed in the text. Hits are considered enriched if they are *≥* 3 fold over the control and have a p-value <0.05. B) GFP vector, CCSer2^WT^, or CCSer2^ΔCC^ were expressed in and co-immunoprecipitated from U2OS cells with anti-FLAG resin. Membranes were probed with α-FLAG, α-dynein heavy chain (DHC), α-Lis1, α-Ndel1, and α-GAPDH antibodies. C) Domain schematics of CCSer2 (green) and Ndel1 (pink) constructs used. Interaction between the two proteins is shown with a colored line above the schematics. D) AlphaFold prediction of CCSer2 and Ndel1 shown in (A) with most disordered regions of CCSer2 removed for clarity. E) Binding curve between Ndel1 (circles) or NT-Ndel1 (triangles) and CCSer2. n = 3 for both samples. Error bars are mean ± SD. F) Fluorescence microscopy image of U2OS cells expressing CCSer2^WT^, stained for α-GFP and α-tubulin. G) The ratio of the intensity around the cell perimeter to the cytoplasmic intensity for CCSer2^WT^ and microtubules. n = 50 cells analyzed. Wilcoxon t-test performed, error bars are mean with standard deviation. H) Temporal hyperstack colored as indicated for overexpressed CCSer2^WT^, CCSer2-SxNN^ALL^, and CCSer2-SxNN^1,4^. I) Domain schematic of CCSer2 showing relative position of S-x-L-P motifs and associated mutants used J) Number of plus-end comet structures in cells expressing CCSer2^WT^, CCSer2-SxNN^ALL^, or CCSer2-SxNN^1,4^. n = 26, 21, 19 cells analyzed for CCSer2^WT^, CCSer2-SxNN^ALL^, or CCSer2-SxNN^1,4^, respectively. Error bars are median ± interquartile range. Statistical analysis performed with a Kruskal-Wallis test with Dunn’s multiple comparisons.

To narrow down the list of potential interactors, we examined hits that were enriched in all three datasets. In addition to Lis1 and Ndel1, there were four other proteins enriched in all datasets (Figure 1A). These hits include the Ndel1 paralogue, Nde1, which can form a complex with Ndel1 and also binds Lis1;^28^ dynein intermediate chain, which is an integral subunit of the dynein motor machinery; TCPD, which is a subunit of the TriC chaperone complex and may facilitate Lis1 β-propellor folding;^29^ and CCSer2, a poorly characterized and largely unstructured protein (Figure S1E, F). CCSer2 was a particularly strong hit with peptides found in all experimental replicates for each dataset, but not in the control. We also identified the CCSer2 homologue CCSer1 in both Ndel1 datasets (Figure 1A). We focused our attention on CCSer2 rather than CCSer1 because it was a robust hit in all datasets and was expressed in all cultured cell lines we tested (Figure S1G). CCSer1, in contrast, was undetectable in most cell culture lines and is generally expressed at a lower level in humans (Figure S1G).^30^

Next, we set out to map the interaction between CCSer2, Ndel1, and Lis1. First, we tested if CCSer2 co-immunoprecipitated Ndel1, Lis1, or dynein in U2OS cells using an N-terminally sfGFP- and C-terminally FLAG-tagged construct of CCSer2 (CCSer2^WT^) (Figure 1B; S1H). While CCSer2^WT^ did co-precipitate Ndel1, we did not detect specific immunoprecipitation of Lis1 or dynein, suggesting that CCSer2 does not bind to either protein directly (Figure 1B; S1H). We next used AlphaFold to predict potential sites of interaction between Ndel1 or Lis1 and CCSer2.^31,32^ Consistent with the co-immunoprecipitation results, no high-confidence interactions were predicted to form between CCSer2 and Lis1. However, AlphaFold predicted a high-confidence interaction between CCSer2 and Ndel1 (Figure S1I-J). The putative interface is comprised of four helix bundle formed from amino acids 657-687 in CCSer2 and amino acids 246-278 in Ndel1 (Figure 1C, D; S1K). In both proteins these regions are predicted to form coiled coils in the absence of any binding partners.

To test the validity of the CCSer2-Ndel1 four-helix bundle predicted by AlphaFold, we generated a CCSer2 construct missing the coiled coil that contains the potential Ndel1 binding site (CCSer2^ΔCC^) (Figure 1C). As predicted, CCSer2^ΔCC^ no longer coimmunoprecipitated Ndel1 (Figure 1B; S1H). Next, we recombinantly expressed and purified a minimal construct of CCSer2 that contains only the Ndel1-binding region (amino acids 650-680) fused to a HaloTag (CCSer2^650–850^) and tested binding to Ndel1 in vitro using a bead-based depletion assay (Figure 1C). We found that the Ndel1-CCSer2^650–850^ interaction was high affinity, with a Kd of *∼*14 nM (Figure 1E). CCSer2^650–850^ did not interact with a truncation of Ndel1 (NT-Ndel1) that has the predicted CCSer2 binding region deleted, which confirms the specificity of the interaction (Figure 1C, E). Additionally, neither Lis1 nor dynein showed any appreciable binding to CCSer2^650–850^ (Figure S1L). Together, these findings are consistent with the AlphaFold prediction and suggests that CCSer2 binds directly to Ndel1’s C-terminal coiled coil. Further, these results suggest that CCSer2 is found within the Lis1 interactome likely due to Lis1-Ndel1 association.

### CCSer2 localizes to the microtubule plus-end and cell periphery

To explore the cellular function of CCSer2, we first determined the localization in human cells. In fixed samples, we observed strong colocalization with microtubules, often accompanied by notable enrichment in CCSer2 signal around the cell perimeter (Figure 1F,G; S2A). When imaged live, as was observed with the mouse homologue, CCSer2^WT^ displayed many dynamic, comet-like structures reminiscent of proteins that track microtubule plus-ends (+Tips) (Figure 1H, Supplemental movie-1).^33^ Consistent with microtubule plus-end localization, CCSer2^WT^ puncta colocalized with the master plus-end regulator, EB1 (Figure S2B). Indeed, CCSer2 contains four EB1-binding S-x-I/L-P motifs, where “x” corresponds to any amino acid (Figure S2C).^34,35^ We found that mutation of all four motifs or just the two most conserved (CCSer2-SXNN^ALL^ and CCSer2-SXNN^1,4^, respectively) significantly reduced plus-end localization of exogenously expressed CCSer2, suggesting that CCSer2’s plus-end localization is EB1 dependent (Figure S1H, I, Supplemental movie-2). CCSer2^ΔCC^ also showed a reduction in comet-like structures, suggesting that the coiled coil that contains the Ndel1 binding site also promotes plus-end localization (Figure 1J, S2D).

### CCSer2 depletion causes cell migration defects in zebrafish primordium

To investigate the role of CCSer2 in vivo, we analyzed CCSer2 expression and function in zebrafish. Zebrafish have two *ccser2* genes, *ccser2a* and *ccser2b*. First, we characterized the expression of *ccser2a* and *ccser2b* at 30 hours post-fertilization (30 hpf) and observed that they both are ubiquitously expressed (Figure S3A) and highly maternally deposited in developing zygotes (Figure S3B). To probe CCSer2 function, we created a *ccser2a;ccser2b* double mutant using CRISPR-Cas9 mutagenesis to disrupt both genes. In *ccser2a;ccser2b* double heterozygous crosses, double mutant offspring survived at sub-mendelian ratios, consistent with the loss of the *ccser2* homologue in mouse litters (Figure S3C).^33^ However, surviving double mutant animals were grossly normal. This bimodal effect, where the double mutant is lethal in most embryos but escapers are healthy, is likely due to variable genetic compensation by *ccser2*’s paralog, *ccser1* because in stable mutant zebrafish lines, paralogs are often upregulated to compensate for genetic deletion.^36^

To explore the function of CCSer2, we silenced both *ccser2* genes in zygotes and embryos using Cas9 and guide RNAs in G0 animals (i.e. crispants), raised them to 4 days post fertilization (dpf) and assessed phenotypes. This crispant approach does not cause upregulation of paralogous genes and is therefore a useful tool to assess phenotypes independent of paralog compensation.^37^ Indeed, we observed that 4 dpf *ccser2a;ccser2b* crispants had overt defects in the posterior lateral line (pLL) structure. The zebrafish pLL is a mechanosensory system composed of sensory neurons and the mechanosensory organs they innervate, called neuromasts. Axons in the pLL of *ccser2a;ccser2b* crispants were significantly truncated and fewer sensory organs were formed compared to uninjected siblings (Figure S3D-F). To confirm this phenotype, we attenuated CCSer2a and CCSer2b protein levels with morpholino antisense oligonucleotides targeting the start codon of *ccser2a* and *ccser2* mRNA. Because morpholinos sterically inhibit protein translation, they are another suitable approach to explore functions of genes that may have compensatory paralogs. *ccser2a;ccser2b* morphants displayed defects in pLL structure that were identical to *ccser2a;ccser2b* crispants (Figure 2A-C). Together, these data demonstrate that loss of CCSer2 in zebrafish causes a specific defect in the formation of the pLL.

**Figure 2.**
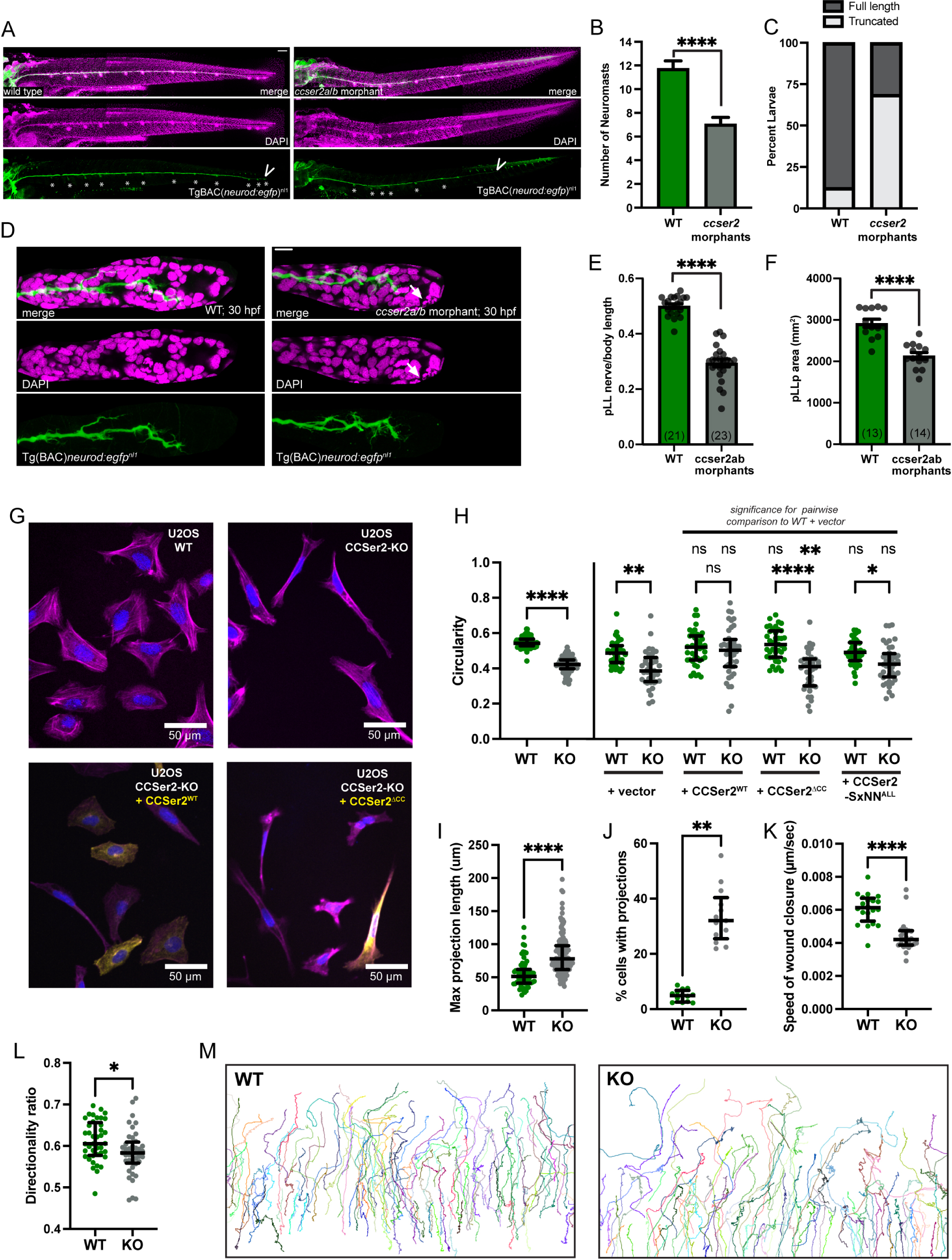
CCSer2 regulates cell migration. A) Fluorescence microscopy image of fixed wild type and *ccser2a/b* zebrafish morphants stained for DAPI (magenta) and neurons (green). Arrow indicates the end of the pLL and asterisks mark neuromasts. B) Number of neuromasts in wild type and *ccser2a/b* morphants as shown in (A). Chi-square; p<0.005) C) Percent of wild type of *ccser2a/b* morphants with truncated lateral lines, as shown in (A). Chi-square; p<0.0001). D) DAPI (magenta) and neurons (green) in single plan image of the pLL primordium at day 30 hpf for wild type embryos or *ccser2a/b* morphants. E) pLL length at 30dpf for wild type or *ccser2a/b* morphants. F) pLL primordium (pLLp) area for wild type or *ccser2a/b* morphants. (ANOVA; p<0.0001). Scale bar = 100 mm in (A); 10 mm in (D) (G) Fluorescence microscopy images of fixed U2OS-WT, CCSer2-KO, and CCSer2-KO with exogenous expression of CCSer2^WT^ or CCSer2^ΔCC^, stained with phalloidin to visualize actin (pink), α-GFP to visualize CCSer2^WT^ and CCSer2^ΔCC^ (yellow), and DAPI to visualize nuclei (blue). H) The average circularity of cells in a field of view for WT and CCSer2-KO cells (left of line). n = 73 fields of view analyzed across three biological replicates for WT and CCSer2-KO cell samples. Error bars are median ± interquartile range. Statistical analysis performed with a Mann-Whitney test. Average circularity of CCSer2-KO cells transfected with vector, CCSer2^WT^, CCSer2^ΔCC^, or CCSer2-SxNN^ALL^ (right of dotted line). N = 40 fields of view analyzed across three biological replicates for CCSer2-KO expressing vector, CCSer2^WT^, CCSer2^ΔCC^, or CCSer2-SxNN^ALL^. Significance determined with a Kruskal-Wallis test with Dunn’s multiple comparisons. Error bars are median ± interquartile. I) Maximum length of each projection (protrusion that is thinner than 10 *µ* M and lasts for at least 18 minutes) formed in WT and CCSer2-KO cells over a period of 24 hours. n = 64 and 127 for WT and CCSer2-KO cells, respectively. J) The percent of WT or CCSer2-KO cells that formed projections within the 20X field of view. n = 13 fields of view analyzed for both WT and CCSer2-KO cells across two biological replicates. K) Speed of wound closure for WT and CCSer2-KO cells. n = 20 and 23 fields of view across three biological replicates. L) The directionality ratio (net displacement/total distance) of individual tracks of WT or CCSer2-KO cells migrating during a wound healing assay. SIR-DNA stained nuclei were used as a fiducial for tracking. n = 20 fields of view analyzed for both WT and CCSer2-KOs across two biological replicates with two technical replicates each. I-L) Error bars are median ± interquartile range. Statistical analysis performed with a Mann-Whitney test. M) Representative migratory tracks of WT and CCSer2-KO nuclei as cells migrate to fill a wound at the top of the image. Each color designates an individual cell trajectory.

The pLL develops early, between 20 and 48 hours post fertilization (hpf), through the migration of the pLL primordium (Figure S3G-J). The primordium undergoes repetitive rounds of cell division in the leading region followed by organ patterning and deposition of neuromasts from the trailing region as it migrates from the embryonic head to tail. As the primordium migrates, it tows the growth cones of the pLL axons (Figure S3G-J). We hypothesized that failed pLL development resulted from defects in pLL primordium migration upon CCSer2 knockdown. To test this, we determined the impact of CCSer2 loss of function on pLL primordium migration in *ccser2a;ccser2b* morphants at 30 hpf, when the primordium should have migrated 50% of the way through the trunk. *ccser2a;ccser2b* morphants displayed significant reduction in pLL primordium migration, as indicated by shorter pLL nerves (Figure 3D, E; S3K). In addition, pLL primordia area was reduced and more apoptotic cells were observed compared to wild type controls (Figure 3D, F; S3L, M). Together, these data implicate CCSer2 in collective migration in vivo.

**Figure 3.**
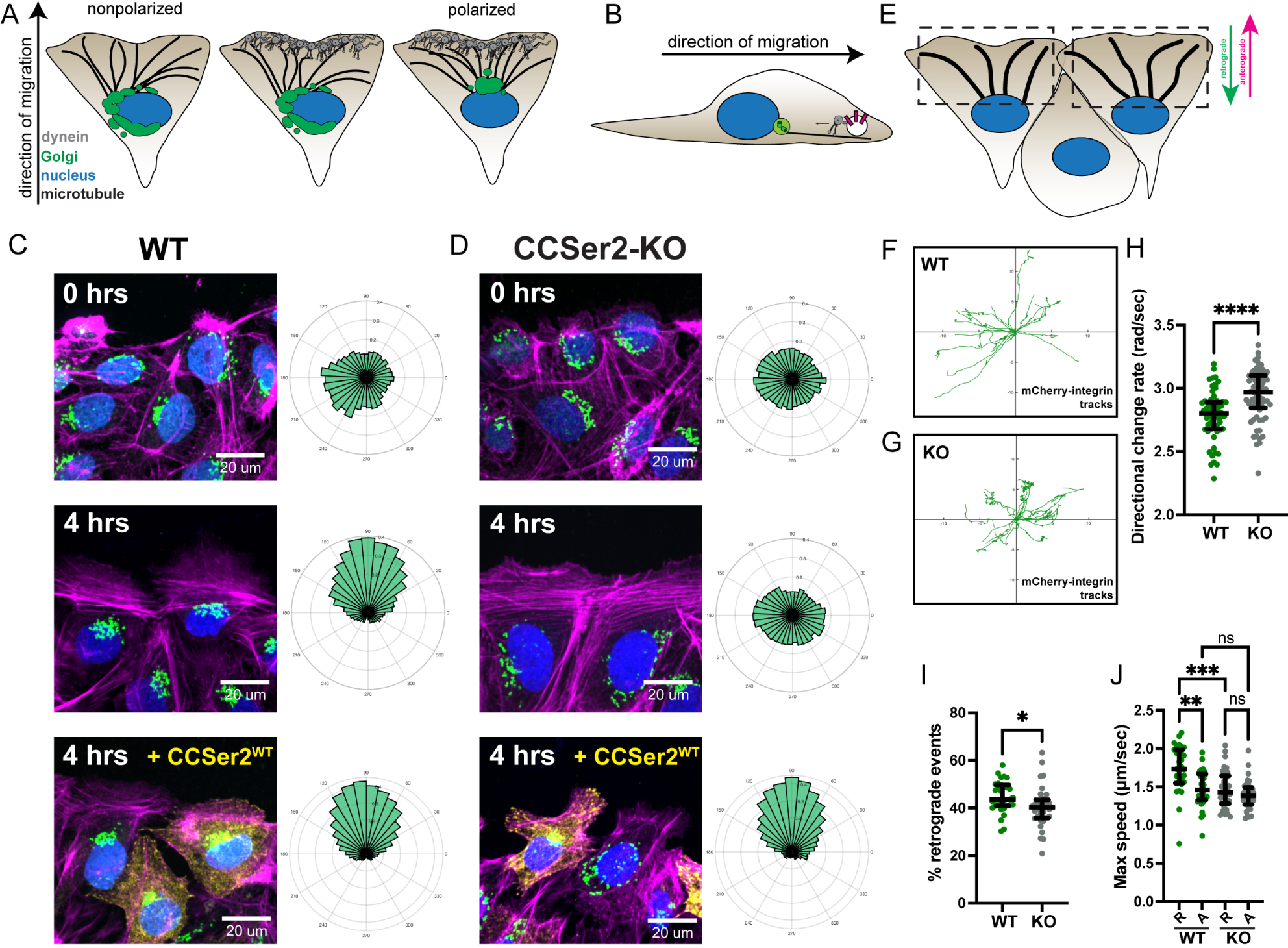
CCSer2 knockout causes a decrease in microtubule polarization during migration and retrograde trafficking defects of integrins. A) Illustration of dynein’s role in microtubule polarization during cell migration. Dynein anchored to the leading edge of a migrating cell facilitates the polarization of the nuclear-centrosomal axis by pulling the on cortical microtubules. The Golgi apparatus is localized near the centrosome and reports on centrosome position. B) Internalized integrins are trafficked by dynein before being shuttled to recycling endosomes or the lysosome. C, D) Fluorescence microscopy images of fixed WT (C) or CCSer2-KO (D) cells at 0 hours and 4 hours post wounding or 4 hours post wounding with expression of exogenous CCSer2^WT^. Cells were stained with α-GM130 (green) to label Golgi, phalloidin (pink) to label actin, DAPI (blue) as to label the nucleus, and α-GFP to label CCSer2^WT^ transfected cells (yellow). Rose plots next to each image indicate the position of the Golgi around the nucleus relative to the leading edge, compiled from at least 86 cells per sample across three biological replicates. E) Schematic of a cell in a confluent layer migrating upwards to fill in a wound. The orientation of the polarized microtubule network within the dashed box establishes upward moving vesicles as anterograde (pink arrow) and downward moving vesicles as retrograde (green arrow). F, G) Representative tracks longer than 5 *µ*m of moving integrins in WT (F) or CCSer2-KO (G) cells. 17 tracks are shown from the WT cell and 14 tracks are shown from the CCSer2-KO cell. H) The directional change rate of both anterograde and retrograde individual integrin tracks averaged per cell. n = 62 and 66 track averages analyzed for 31 and 33 WT and CCSer2-KO cells, respectively, across three biological replicates. I) Percentage of retrograde events in WT and CCSer2 KO cells. n = 31 and 33 cells analyzed for WT and CCSer2-KO cells, respectively, across three biological replicates. H, I) Error bars are median ± interquartile range. Statistical analysis performed with a Mann-Whitney test. J) The maximum speed of the averaged retrograde and anterograde integrin tracks per cell of WT and CCSer2-KO cells. n = 31 and 33 cells analyzed for WT and CCSer2-KO cells, respectively, across three biological replicates. Error bars are median ± interquartile range. Statistical analysis performed with a Kruskal-Wallis test with Dunn’s multiple comparisons.

### CCSer2 deletion causes a reduction in directional persistence during migration

To determine the cellular defect that gave rise to failed migration in vivo, we used CRISPR-Cas9 to delete CCSer2 from U2OS cells (CCSer2-KO) (Figure S4A, B). CCSer2-KOs showed moderately slower growth compared to control cells, which is likely because of a mild delay in anaphase onset (Figure S4C). However, the most noticeable phenotype was that CCSer2-KO cells had a striking, hyper-elongated cell shape, resulting in a significant reduction in circularity as compared to control cells (Figure 2G, H).

To understand the basis for the change in cell shape, we imaged wild type (WT) control and CCSer2-KO cells live for 24 hours. For control cells, we observed mesenchymal-style migration, with a large flat lamellipodium at the leading edge and a short, retracting tail at the back of the cell (Supplemental movie-3, 4). CCSer2-KO cells, in contrast, displayed highly aberrant migratory behavior, which is consistent with the migratory defect observed in zebrafish primordium (Supplemental movie-5, 6). The KO cells did not appear to form lamellipodia and instead generated long, filopodium-like projections that routinely exceeded 100 µm in length and were almost never observed in control cells (Figure 2I, J). The projections were often dynamic and seemed to actively extend away from the cell body or failed to retract as the cell moved. The propensity for the CCSer2-KO to form projections explained the elongated cell morphology we observed in the fixed samples, as we could directly observe the cells elongate as they failed to retract the projections during migration (Supplemental movie-6). We also observed that the projections would sometimes snap back rapidly, resulting in a cell that appeared rounder and more akin to control cell shapes (Supplemental movie-6). The aberrant morphology did not affect migration rates as the KO cells migrated at the same speed as control cells (Figure S4D). To probe directional migration, we utilized a wound-closure assay and found that CCSer2-KO cells had reduced directional persistence, resulting in slower speed of wound closure (Figure 2K-M; Supplemental movie-7, 8). Together, these results suggest that CCSer2 deletion results in aberrant cell morphology during migration and causes a decrease in directional persistence.

To verify that CCSer2 deletion was the cause of the observed migration defects, we carried out rescue experiments and measured cell shape in fixed samples as a proxy for the migration defects we observed live. Indeed, exogenous expression of CCSer2^WT^ but not vector alone, rescued the elongation defect in CCSer2-KO cells and resulted in a circularity comparable to control (Figure 2G, H; S4E). We next transfected CCSer2-KOs with CCSer2^ΔCC^ or CCSer2-SxNN^ALL^ to determine if CCSer2’s ability to bind Ndel1 or EB1 is important for cell migration. CCSer2^ΔCC^ was unable to rescue the cell shape defect caused by CCSer2 deletion, suggesting that the coiled coil containing the Ndel1 interaction site is important for CCSer2 function during migration (Figure 2G, H). In contrast, CCSer2-SxNN^ALL^ was able to partially rescue the elongated shape in CCSer2-KO cells (Figure 2H; S4E). This result indicates that impairment of CCSer2’s plus-end localization does not severely abrogate its function and suggests that CCSer2 does not operate from the microtubule plus-end to support cell migration.

### CCSer2 knockout causes a defect in microtubule polarization and integrin trafficking by decreasing dynein localization at the leading-edge

Since we found that CCSer2 is a Ndel1 binding protein and CCSer2 depletion causes cell migration defects in zebrafish and U2OS cells, we hypothesized that CCSer2 may function to regulate dynein activity during migration. While migration is largely an actin driven process, microtubules and microtubule motors are involved. Dynein has two main roles during cell migration. First, in many cell types, dynein helps to establish the nuclear-centrosomal axis.^38^ Here, populations of dynein and dynactin are recruited to cell-cell contacts and the leading-edge of migratory cells where they pull on cortical microtubules to reposition the centrosome such that it is between the leading edge and the nucleus (Figure 3A).^39–42^ During this process, dynein also contributes to rotational movement of the nuclei.^42^ Centrosome repositioning ensures that the microtubule network becomes polarized and is aligned with the vector of migration (Figure 3A). A polarized microtubule network is essential for persistent directional movement because it allows for efficient trafficking of signaling proteins and focal adhesion components to and from the leading edge.^38^ Disruption of this process leads to a reduction in directional persistence during migration. Dynein’s second function during migration is to traffic cargo, including endocytosed focal adhesion components, away from the cell periphery (Figure 3B).^43^

The reduction in directional persistence observed in the CCSer2-KO cells is consistent with impaired dynein function during cell migration (Figure 2L, M). Therefore, we set out to determine if CCSer2 depletion affects migration-specific dynein functions. First, we tested if CCSer2 deletion affects centrosome positioning during migration. To test this, we monitored confluent sheets of cells as they migrated to fill a wound. We stained the Golgi network as a reporter for microtubule polarization because its position relative to the nucleus is easy to visualize and because it becomes repositioned with the centrosome. Immediately after generating the wound, the probability of finding Golgi was equal at all angles around the nucleus in both control and CCSer2-KO cells, which is indicative of a non-polarized microtubule network (Figure 3C, D, top). After four hours, all control cells displayed a polarized microtubule network, with the Golgi positioned immediately in front of the nucleus and oriented toward the leading edge (Figure 3C, middle). In contrast, CCSer2-KO cells rarely achieved a polarized microtubule network, with the Golgi network remaining radially distributed around the nucleus (Figure 3D, middle). Exogenous expression of CCSer2^WT^ rescued the polarization defect and generated Golgi positioning profiles in CCSer2-KO cells that were indistinguishable from control samples (Figure 3C, D, bottom).

In addition to establishing the nuclear-centrosomal axis, dynein supports the retrograde trafficking of endocytosed integrins during migration (Figure 3B). To test if CCSer2-KO affects integrin trafficking, we imaged the intracellular movement of exogenously expressed mCherry-α5-integrin-12 in control and CCSer2-KO cells. To discriminate between retrograde and anterograde events, we imaged confluent cells as they migrated to fill a wound and limited our analysis to events that occurred between the leading edge of the cell and the nucleus (Figure 3E). Events that showed a net movement away from the leading edge were considered retrograde, while events that had net movement away from the nucleus were anterograde (Figure 3E). While there was no discernable difference in the net displacement of mCherry-α5-integrin-12 puncta in either cell type, puncta in CCSer2-KO cells displayed far more bidirectional movement than in control cells (Figure 3F-H; S5A). Consistent with a retrograde-specific trafficking defect, CCSer2-KO cells had a lower percentage of retrograde events, and the maximum speed (but not the mean speed) of retrograde events was slower than in control cells (Figure 3I, J; S5B). There was no discernable difference between the maximum speed of anterograde events in control and CCSer2-KO cells (Figure 3J). Altogether, these data suggest that CCSer2 deletion hinders integrin trafficking, with a bigger negative impact on retrograde events. Together, these results support a role for CCSer2 in migration-specific dynein functions and explain the defect in cell migration observed in CCSer2-KO cells.

### CCSer2 exclusively regulates the retrograde trafficking of cargos that require cortically localized dynein

To test if the trafficking defect we observed in CCSer2-KO cells was specific to centrosomes and integrin cargo during migration, or if it applied to all endosomal cargo regardless of the migratory state of the cell, we labeled early endosomes with an antibody against EEA-1 and determined the radial distribution of endosomes with respect to the center of the nucleus in control and CCSer2-KO cells. We conducted this analysis with cells plated at a low density (to ensure cells were not undergoing collective migration) and we excluded hyper-elongated KO cells. In control cells, a large proportion of the EEA-1-positive vesicles were clustered closely around the nucleus (Figure 4A, B). Consistent with a retrograde-trafficking defect, we observed that EEA-1 was significantly dispersed in CCSer2-KO cells (Figure 4A, B). The mean intensity of EEA-1 staining per cell was not significantly different between control and CCSer2-KO cells (Figure S6A), which indicates that there is not a difference in the number of EEA-1 positive vesicles (i.e. not an endocytosis defect), but a defect in the trafficking steps that occur downstream of internalization. Importantly, expression of CCSer2^WT^, but not vector or CCSer2^ΔCC^ rescued the EEA-1 mislocalization in CCSer2-KO cells (Figure 4C-E; S6B), suggesting that the defect is caused by CCSer2 deletion and that the coiled coil containing the Ndel1 binding site is required for rescue.

**Figure 4.**
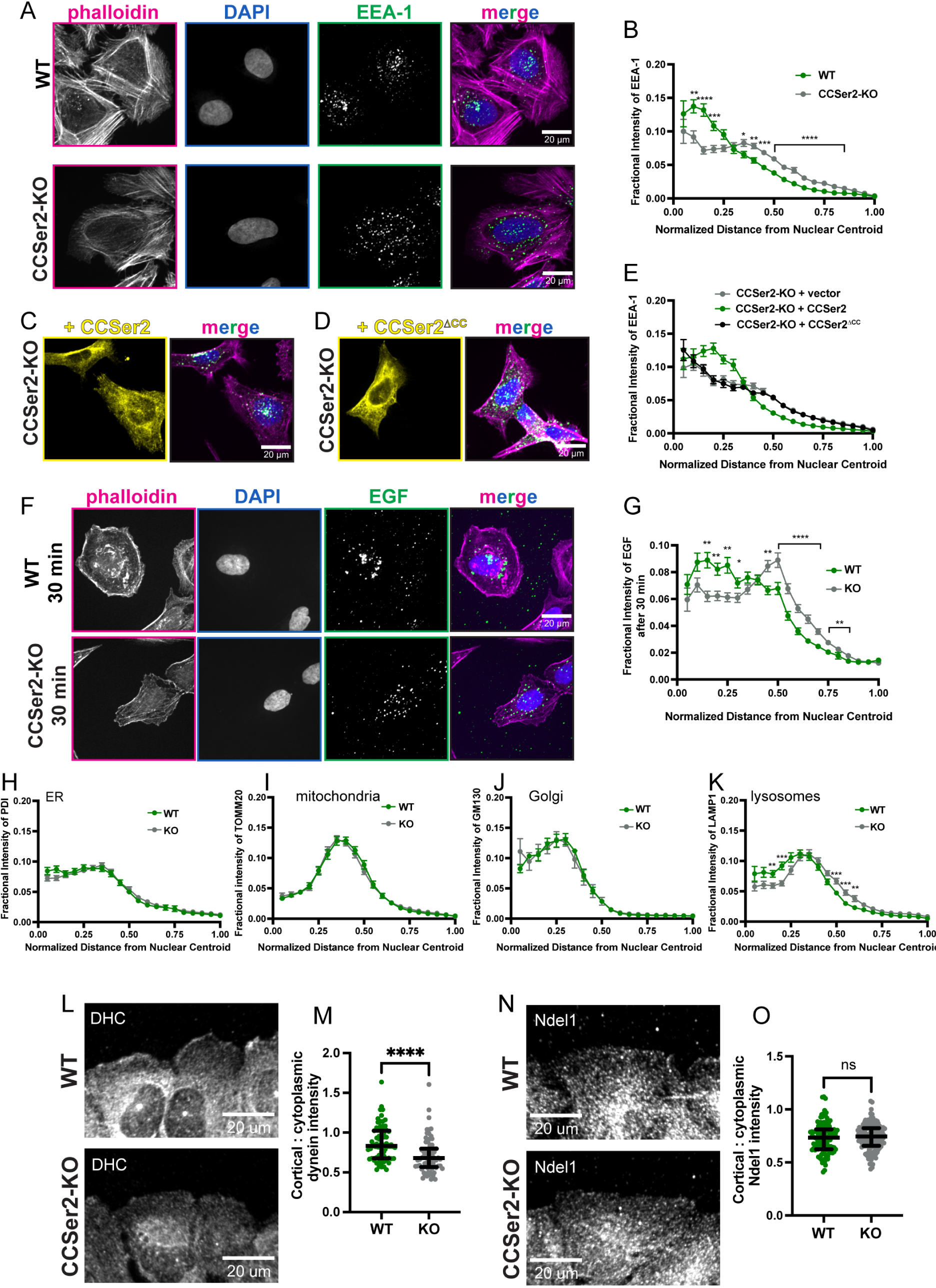
CCSer2 supports the retrograde trafficking of early endosomes and promotes dynein localization to the cell periphery. A) Fluorescence microscopy images of fixed WT and CCSer2-KO cells stained with phalloidin to visualize actin (magenta), α-EEA-1 to visualize early endosomes (green), and DAPI to visualize nuclei (blue). B) Fractional distribution of EEA-1 endosomes from the centroid of the nucleus to the cell edge. n = 37 cells analyzed for both samples across three biological replicates. C, D) Fluorescence microscopy images of fixed CCSer2-KO cells transfected with CCSer2^WT^ (C) or CCSer2^ΔCC^ (D) and stained with phalloidin to visualize actin (magenta), α-EEA-1 to visualize early endosomes (green), α-GFP to visualize CCSer2^WT^ or CCSer2^ΔCC^ (yellow), and DAPI to visualize nuclei (blue). E) Fractional distribution of EEA-1 endosomes from the centroid of the nucleus to the cell edge for CCSer2-KO cells transfected with vector, CCSer2^WT^, or CCSer2^ΔCC^. n = 44 cells analyzed per sample across three biological replicates. F) Fluorescence microscopy images of fixed WT and CCSer2-KO cells treated with EGF-555, shown in green. Cells were stained with phalloidin to visualize actin (magenta) and DAPI to visualize nuclei (blue). G) Fractional distribution of EGF-555 intensity from the centroid of the nucleus to the cell periphery in WT and CCSer2-KOs. n = 89 and 85 cells analyzed for WT and CCSer2-KOs, respectively, across three biological replicates. Error bars shown are mean ± SEM. H-K) Fractional distribution from nuclear centroid to cell edge for ER intensity (stained with α-PDI) (H), mitochondria (stained with α-TOMM20) (I), Golgi apparatus (stained with α-GM130) (J), and lysosomes (stained with α-LAMP1) (K) in WT or CCSer2-KO cells. n= 34 cells for ER, 35 for mitochondria, 37 for Golgi, and 37 for lysosomes, obtained from three biological replicates. B, E, G, and H-K) Error bars shown are mean ± SEM. Statistical analysis performed with multiple, Mann-Whitney tests and a False Discovery Rate (FDR) of 1%. L, N) Fluorescence microscopy images of WT and CCSer2 KO cells fixed 30 minutes post wounding and stained with α-dynein heavy chain (α-DHC) (L) or α-Ndel1 (N). The wound is positioned so that the cells are migrating towards the top of the image. M, O) Quantification of dynein (M) or Ndel1 (O) localization at the leading edge. The intensity of dynein or Ndel1 signal in one-micron band at the leading edge was divided by the mean intensity 6.5 microns into the cell. n = 75 cells analyzed for both WT and CCSer2-KOs across three biological replicates. Error bars represent median with interquartile range. Significance determined with a Mann-Whitney test.

The cells visualized in this assay may still be undergoing some focal adhesion remodeling and thus a proportion of endosomes still may contain integrins. Therefore, it is possible that the EEA-1 localization defect we observe is caused by specific mislocalization of focal adhesion derived endosomes, and not all endosomes. To test this, we monitored the localization of Epidermal Growth Factor (EGF), which is an endocytosed signaling molecule whose internalization is independent from focal adhesion turnover. Here, we bathed cells in fluorescently tagged EGF and monitored localization at 5 minutes and 30 minutes. There was no difference in the distribution of EGF in either cell type 5 minutes after internalization (Figure S6C, D). However, after 30 minutes, the control cells showed most internalized EGF clustered near the nucleus (Figure 4F, G). In contrast, the CCSer2-KO cells displayed more dispersed EGF puncta, which is consistent with a dynein trafficking defect (Figure 4F, G). Together, these results suggest that CCSer2-KO cells have impaired dynein-driven retrograde trafficking of all early endosomes. Further, these findings suggest that the integrin trafficking defect observed in CCSer2-KO is likely an outcome of impaired endosome trafficking and not specific to integrin cargo.

Dynein is a multifunctional motor. In addition to trafficking endosomes, dynein supports the retrograde motility of hundreds of other cargos. To test if CCSer2-KO affects all cargo that moves in the retrograde direction, we looked at the cellular distribution of four additional model cargo (endoplasmic reticulum (ER), mitochondria, Golgi, and lysosomes) using the same methodology that we used to measure endosome distribution. If CCSer2 regulates all retrograde trafficking events, we anticipated that we would observe mislocalization of all cargo. However, if CCSer2 activity specifically affects endosomes, we reasoned that we would only observe mislocalization of lysosomes, which mature from populations of early endosomes. Consistent with the latter hypothesis, we observed that ER, mitochondria, and Golgi all showed identical cellular distribution profiles in both control and CCSer2-KO cells, while lysosomes were moderately dispersed in CCSer2-KO cells (Figure 4H-K; S6E-H).

Given CCSer2’s localization to the microtubule plus-end and the cell periphery, we next asked if CCSer2 controls dynein localization to both cellular regions. CCSer2 depletion had no effect on dynein intensity at microtubule plus-ends (Figure S6I). In contrast, we observed very little dynein signal at the leading edge of migrating CCSer2-KO cells (Figure 4L, M; S6J), suggesting that CCSer2 can promote dynein localization to the cell periphery. Interestingly, CCSer2-KO cells did not change the relative localization of Ndel1 at the leading edge, which suggests that CCSer2 does not promote dynein localization at the cell periphery simply by binding Ndel1-dynein complexes (Figure 4N, O; S6K). These data also explain how CCSer2 deletion only affects dynein-mediated centrosome positioning and endosome trafficking: both cargos require dynein localized to the cell periphery for proper trafficking.

### CCSer2 attenuates the Ndel1-mediated inhibition of dynein activation

We showed that 1) CCSer2 binds directly to Ndel1, but not dynein (Figure 1B-E; S1L) and 2) CCSer2 promotes dynein, but not Ndel1, localization at the cell periphery (Figure 4L-O). At first glance, these two observations are hard to reconcile. However, Ndel1 directly regulates dynein activity by competing with dynactin for dynein binding, thus attenuating formation of the activated transport complex. We reasoned that if CCSer2 inhibits Ndel1 regulation of dynein it would act as a dynein localization factor by promoting the formation of active transport complexes on adaptor-bound cargos near the cell periphery (Figure 5A). To test this hypothesis, we first reconstituted active transport complexes in vitro using purified dynein, dynactin, and the adaptor BicD2 (Figure 5B, C; S7A, B). Here we used a well-characterized truncation of BicD2 that relieves auto-inhibition.^17^ We next confirmed that Lis1 and Ndel1 behaved as we and others have reported. Briefly, incubation with Lis1 caused an increase in the number of processive dynein-dynactin-BicD2 complexes (i.e. landing rate), while inclusion of Ndel1 with Lis1 resulted in a decreased landing rate, compared to Lis1 alone (Figure 5B, C).^17–20,44^

**Figure 5.**
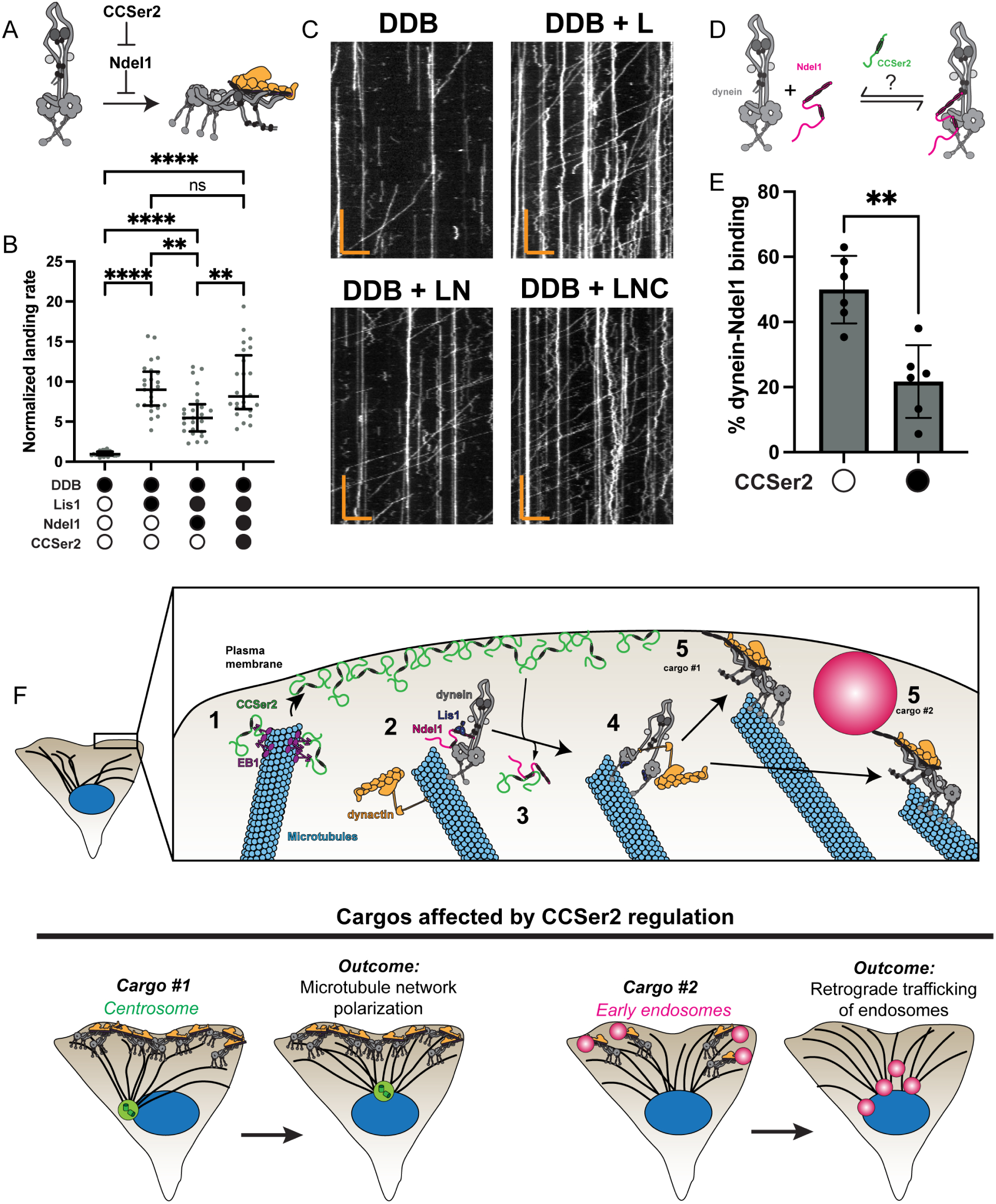
CCSer2 activates dynein via Ndel1 inhibition. A) Schematic of how CCSer2 inhibition of Ndel1 could result in dynein activation. B) Number of processive events per µm of microtubule per nM of dynein for dynein-dynactin-BicD2 (DDB) complexes in the absence (white circles) or presence (black circles) of 50 nM Lis1, 10 nM Ndel1, and/or 80 nM Halo-CCSer2^650–850^. Data are normalized to the dynein-dynactin-BicD2 sample for each replicate. n = 25 microtubules analyzed for each condition. Statistical analysis performed with a Brown-Forsythe and Welch ANOVA with Dunnet’s T3 multiple comparison test. C) Example kymographs of dynein-dynactin-BicD (DDB) in the presence of Lis1 (DDB + L), Lis1 and Ndel1 (DDB + LN), or Lis1, Ndel1, and Halo-CCSer2^650–850^ (DDB + LNC). Y-scale bar is 15 seconds, X-scale bar is 5 µm. D) Schematic of assay to determine if CCSer2^650–850^ affects the dynein-Ndel1 interaction. E) Percent of dynein bound to Ndel1-conjugated beads in the absence (white circle) or presence (black circle) of CCSer2^650–850^. n = 6. Statistical analysis performed with a Mann Whitney test F) Model for CCSer2 delivery to the cortex and spatial regulation of dynein. Step 1) CCSer2 binds to EB1 to associate with growing microtubule plus-ends to reach the cell periphery. Step 2) Dynein-Lis1-Ndel1 form a primed structure that is poised for activation. We have shown this on microtubules, but it is also possible that this structure does not form on the microtubule. Step 3) Ndel1-CCSer2 interaction reduces Ndel1’s affinity for dynein. Step 4) Ndel1 release of dynein allows Lis1-mediated activation of dynein and dynactin binding. Step 5) The fully activated transport complex of dynein-dynactin-adaptor forms with adaptors that are already associated with cellular structures where active dynein will be recruited. Cargo #1) Dynein-dynactin-adaptor complexes form on the actin cortex (adaptor unknown) to reposition the centrosome during migration. Cargo #2) Dynein-dynactin-adaptor complexes form on early endosomes (likely Hook1 and Hook3) to drive retrograde trafficking.

Next, we asked if CCSer2^650–850^ had any impact on the combined effect that Lis1 and Ndel1 had on the landing rate of dynein-dynactin-BicD2 complexes. Remarkably, we observed that inclusion of CCSer2^650–850^ returned the landing rate of dynein-dynactin-BicD2 complexes to the levels observed only with Lis1 present (Figure 5B, C). Importantly, CCSer2^650–850^ had no effect on dynein motility in the absence of Ndel1 (Figure S7C-K). This result indicates that CCSer2 attenuates the inhibition of dynein-dynactin-BicD2 landing rate caused by Ndel1 by inhibited Ndel1-dynein association and suggests that the dynein populations localized via CCSer2 are bound to dynactin and cargo adaptors.

Ndel1 opposes dynein activation in two ways. First, Ndel1 competes with the dynactin subunit p150 for binding to dynein intermediate chain. In this way, Ndel1 inhibits the association of dynein and dynactin. Second, Ndel1 competes with dynein for Lis1 binding and thus prevents Lis1-mediated activation of dynein. To determine how CCSer2 attenuates Ndel1 regulation of dynein, we asked if CCSer2^650–850^ affected dynein-Ndel1 or Lis1-Ndel1 binding using a bead-based depletion assay with purified components (Figure 5D; S7L). Consistent with our model, we observed that dynein-Ndel1 binding was reduced in the presence of CCSer2^650–850^ (Figure 5E). Lis1-Ndel1 interaction was not affected by CCSer2^650–850^ (Figure S7M). These result suggests that CCSer2 and dynein compete for Ndel1 interaction and is consistent with previous reports that Ndel1’s C-terminal coiled coil that binds to CCSer2 (Figure 1D, E; S1I-K) also mediates dynein binding.^21,45^ Altogether, our data shows that CCSer2 promotes dynein localization at the leading edge by directly modulating Ndel1’s effect on the dynein activation equilibrium.

## Discussion

We set out to discover novel regulatory circuits that gate dynein activity in cellular space. We reasoned that the proteins Lis1 and Ndel1, which are key regulators of dynein localization and activation, may be nodes through which spatial-type information is conveyed to dynein. We used proximity dependent biotinylation coupled to MS to identify proteins that are positioned to modulate Lis1 and Ndel1 activity. We identified CCSer2, a poorly characterized protein that is in the interactome of both proteins. We found that CCSer2 supports dynein localization at the cell periphery to facilitate dynein-mediated centrosome positioning during cell migration and early endosome trafficking. Disruption of these processes via CCSer2 depletion impairs migration in cell culture models and developing zebrafish. Finally, we showed that CCSer2 binds to Ndel1 directly to reduce the inhibitory effect that Ndel1 has on dynein-dynactin-adaptor formation, thus promoting dynein activation and stabilizing populations of dynein on cargo that reside near the cell edge. CCSer2 activity represents the first described dynein regulator that activates dynein in a specific cellular space, rather than on a specific cargo.

Our work describing CCSer2 function also explains how Lis1 and Ndel1 activity can be modulated to promote dynein localization and activation. In our model, by functioning as an inhibitor of Ndel1, CCSer2 effectively “deploys” dynein at the cell cortex, simultaneously promoting its localization and interaction with dynactin and adaptor (Figure 5F). Below, we describe our model of how CCSer2 regulates the dynein transport machinery at the cell periphery, as well as highlight outstanding questions.

*Step 1: CCSer2 associates with microtubules-plus ends to reach the cell periphery*. We showed that CCSer2 localizes to the cell periphery (Figure 1F, G). We speculate that CCSer2 localizes by binding to actin or actin-associated proteins at the cell cortex, however more work is required to test this. CCSer2 also is a microtubule-plus end tracking protein (Figure 1H-J). We hypothesize that EB1-driven microtubule-plus end localization facilitates the efficient delivery of CCSer2 to the cell edge, as has been hypothesized for other cortical proteins (Figure 5F, Step 1).^46,47^ However, it is important to emphasize that CCSer2 mutants that can’t bind EB1 and have reduced colocalization to the plus-end still rescue migration defects in CCSer2-KO cells (Figure 2H; S4E). This suggests that either overexpression of CCSer2 bypasses the need for plus-end delivery to the cortex or that microtubule association of CCSer2 is not central to its function.

*Step 2: Dynein-Lis1 complexes are held in a “primed” conformation by Ndel1*. Previous work has shown that, Lis1 and Ndel1 enable dynein’s microtubule plus-end localization.^11,48–50^ Dynactin also localizes to microtubule plus-ends and may promote localization of dynein.^51^ Interestingly, a recent study has revealed that Lis1 and dynein are trafficked to the plus-end via a different mechanism than dynactin and Ndel1, which may highlight the need to prevent aberrant, mislocalized dynein activation.^52^ Once at the plus-end, we speculate that a dynein-Lis1-Ndel1 complex is formed, where Lis1 and dynein are scaffolded together by Ndel1, yet held in an inactive state (Figure 5F, Step 2). We and others have shown that while Ndel1 tethers Lis1 to dynein, it does so in a manner where dynactin can’t bind dynein and Lis1 can’t bind to dynein’s motor domain and is thus inactive.^20,21^ We reason that this state represents a “primed” conformation, which allows multiple dynein motors and the machinery required to activate dynein motility to be recruited to a specific cellular location without forming activated complexes. In this way, dynein can be accumulated in a specific cellular space before Ndel1 release triggers dynein activation. It is important to note that others have observed that in vitro, the Ndel1 homologue, Nde1, facilitates Lis1-mediated activation of dynein rather than inhibits it.^23^ To determine if Ndel1 is also capable of functioning in this way requires further exploration and is not mutually exclusive with our proposed model.

*Steps 3 - 5: CCSer2 disfavors Ndel1-inibition of dynein, thus facilitating dynein activation and recruitment to the cell periphery*. We found that CCSer2 promotes dynein, but not Ndel1, localization to the leading edge (Figure 4L-O). How does CCSer2 accomplish this? We showed that CCSer2 binds directly to Ndel1 but does not bind to dynein (Figure 1B, E). Our in vitro motility experiments suggest that CCSer2 likely functions as an “off-loading factor” that promotes dynein-cortex localization by destabilizing Ndel1 inhibition of dynein (Figure 5F, Step 3). We predict Ndel1-CCSer2 interaction causes Ndel1 to unbind from dynein, which accomplishes two major outcomes (Figure 5F, Step 4): 1) Lis1 that is not Ndel1 bound is free to promote the open conformation of dynein and recruit dynactin. 2) Dynein intermediate chain, which was previously bound to Ndel1, is free to bind to the p150 subunit of dynactin. Both outcomes drive dynein-dynactin-adaptor association and, thus dynein activation (Figure 5F, Step 5). While we are showing these steps as distinct in this model, it is possible that these steps occur simultaneously. More work will be required to determine the order of how dynein, dynactin, and adaptor proteins assemble into complexes.

How does CCSer2 reduce the dynein-Ndel1 binding affinity? We have mapped the region of Ndel1 required to bind CCSer2 to the C-terminal half of Ndel1 and high confidence AlphaFold predictions support these findings (Figure 1D, E; S1J-K). Previously, we found that Ndel1’s C-terminal coiled coil contributes to dynein binding and that by removing it, Ndel1-dynein affinity is significantly reduced.^21^ We hypothesize that CCSer2-Ndel1 binding reduces dynein-Ndel1 interaction by simply sequestering Ndel1’s C-terminal coiled coil away from dynein. Interestingly, the C-terminal coiled coil of Ndel1 is predicted to bind to multiple other proteins that localize at different regions throughout the cell and locally modulate Ndel1 activity.^8,27^ For example, ankyrin-G recruits Ndel1 to the axon initial segment, where Ndel1 plays a role in regulating dynein’s ability to sort somatodendritic cargo, while the interaction between DISC-1 and Ndel1 on mitochondria effects mitochondrial trafficking.^53,54^ Interestingly, both ankyrin and DISC-1 bind Ndel1 in a manner that is reminiscent of how CCSer2 binds, forming helical bundles with Ndel1’s C-terminal coiled coil.^55,56^ We propose that these and other proteins likely function as CCSer2 does and bind to the C-terminal coiled coil of Ndel1 to modulate inhibition of dynein activation in specific cellular locations.

In our model, we have indicated a generic “adaptor” because, in this study, we did not explicitly probe the adaptors that are subject to CCSer2 regulation. Because Ndel1 operates upstream of dynein-dynactin-adaptor binding, CCSer2 regulation of dynein is almost certainly indifferent to the identity of the adaptor. By localizing to the cell cortex, CCSer2’s activity is spatially restricted and thus, only modulates dynein’s access to the subset of cargos that are operated on or originate from the cell periphery. Our model predicts that any adaptor that localizes along the cell edge will experience reduced binding to dynein upon CCSer2 depletion. Further discussion of adaptors involved in the processes described in this study is included below.

*Outcome 1: Dynein activated at the cell cortex during migration helps to position the centrosome and results in a polarized microtubule network*. We showed that CCSer2 promotes dynein localization to the leading edge of migrating cells (Figure 5F, G). Without CCSer2, dynein does not reach the leading edge, cannot position the centrosome, and therefore fails to promote polarization of the microtubule network, resulting in a decrease in directional persistence of migrating cells (Figure 2L, M; 3C, D; 4L-M). This may contribute to the migration defects observed in the pLL primordium cells when CCSer2 is depleted in zebrafish embryos (Figure 2A-F). However, some cells in the pLL position the centrosome posterior to the nucleus during migration, which may not require dynein.^38,57^ More work is required to determine the contribution that dynein and CCSer2 have on establishing the centrosome-nucleus axis in pLL cells.

The adaptor required for centrosome positioning during cell migration is not known, however Par3 is required for dynein activity during this process.^39^ Indeed, Par3 co-immunoprecipates dynein and dynactin subunits and promotes their localization to the leading edge.^39^ Par3 is unlikely to act as a dynein adaptor, as it contains none of the structural hallmarks of any currently identified adaptor. We speculate that Par3 associates with the dynein adaptor responsible for activation during this process because the co-localization of Par3-dynein at the leading edge is dependent on the presence of the light intermediate chain subunit of dynein, which is a key mediator of dynein-adaptor binding.^39^ More work is required to identify the adaptor required for dynein activation and to establish the interaction network of the dynein transport machinery during cell migration.

*Outcome 2: Dynein activation at the cell cortex promotes loading onto and trafficking of early endosomes*. We also found that early endosomes display a retrograde trafficking defect upon CCSer2 deletion (Figure 4A, B). Because early endosomes develop from the plasma membrane, the subpopulation of dynein localized on the cortex is certainly utilized for the early microtubule-dependent retrograde trafficking of these vesicles. The defects in integrin trafficking observed in the CCSer2-KO cells is likely caused by the impaired retrograde trafficking of endosomes generally and is not specific to integrins. It is also possible that a reduction in dynein-driven retrograde trafficking of early endosomes contributes to polarization defects we observed, as intact retrograde trafficking is required to support cell polarization.^43,58^ Disruption of the movement of endosome cargo likely contributes to the migration defects observed in zebrafish (Figure 2A-F). Hook1 and Hook3 are the adaptors implicated in early endosome trafficking.^59^ Indeed, Hook3 shows striking localization to the cell-periphery, which is consistent with its role in dynein-driven endosome trafficking and suggest that its interaction with dynein is likely affected by CCSer2.^60^

*CCSer2 activity represents a new lens through which to examine dynein regulation*. The function of CCSer2 that we have described here not only reveals how Ndel1 can support dynein localization and activation, but it provides a new archetype for how ubiquitously expressed motor machinery can be activated with spatial specificity. CCSer2, by inhibiting Ndel1 inhibition exclusively at the cell periphery, functions as a localized activator of dynein motility. The logic that governs the CCSer2-Ndel1-dynein triad is found in multiple biological processes. For example, nearly every kinase studied has regulators that activate via disinhibition of inhibitors and the outcome of this kind of multi-tiered regulatory network is fine-grained, spatially controlled phosphorylation of target protein substrates. To our knowledge, our work in this study is the first example of such a regulatory circuit that controls the activity of a motor protein. Further, this finding offers insight into a long-standing question in the field of dynein research: how can a single dynein motor traffic all retrograde moving cargo with spatial and temporal precision? We suggest that precise activation of dynein motility is achieved by regulation in multiple dimensions, where the localization of dynein adaptors and proteins that disinhibit Ndel1 contribute to tunable dynein activation on specific cargo in discrete cellular microenvironments.

## Author Contributions

M.E.D and J.L.Z. conceived of and planned the project. J.L.Z., D.G., and A.M.Z. generated all reagents, performed all experiments, and analyzed all data using U2OS cells. W.S. and J.P.G. generated all reagents, performed all experiments, and analyzed all data using purified components in vitro. C.S. and C.M.D. generated all reagents, performed all experiments, and analyzed all data involving zebrafish. M.E.D. wrote the manuscript with input from all authors.

## Acknowledgements

The authors thank Drs. Kristen Verhey, Ryoma Ohi, David Sept, Michael Cianfrocco, Eleanor Clowney, Jayakrishnan Nandakumar, Richard Baker, John Salogiannis, Richard McKenney, and William Redwine as well as members of the DeSantis, Cianfrocco, Ohi, Verhey and Sept labs for helpful discussions and/or feedback on the manuscript. We are especially grateful to Dr. Samara Reck-Peterson for resources and generous support for reagent generation. We thank Dr. Zaw Min Htet, Ian Hollyer, and Tien Phuoc Tran for help with the BioID experiments. We are very grateful to Dr. Monika Dzieciatkowska and the University of Colorado Mass Spectrometry Facility for BioID data acquisition and analysis. This work was supported by NIH-R35GM146739, NSF-2142670, and NIH-R00GM127757 (to M.E.D), NIH-T32 GM145304 (to J.L.Z.).

## Methods

### Cloning and plasmid construction

All constructs were generated with isothermal assembly as described.^61^ The constructs used in this study can be found in Table 1.

### Cell culture and transfection

All cell lines were cultured with 1X DMEM (Corning) with 10% FBS (Gibco), and 1% Penicillin/ Streptomycin (Gibco) at 37°C with 5% CO2. Stable BioID fusion cell lines were cultured with 50 µg/mL Hygromycin B. Stable HEK293 cells expressing Halo-tagged p62 subunit of dynactin were cultured with 50 µg/mL Hygromycin B. The cells were passaged every 3-4 days using 0.25% Trypsin-EDTA (Fisher Scientific) incubated at 37°C for 3-5 minutes to detach cells. All cell lines were tested for mycoplasma contamination using the MycoAlert (Lonza) detection kit every 3-6 months. For transfections, 60,000 - 120,000 cells were plated per well of a 6-well tissue culture treated plate (Fisherbrand) and allowed to adhere overnight. The cells were then transfected with either Lipofectamine 2000 (Thermo Fisher Scientific), Lipofectamine LTX and PLUS reagents (Thermo Fisher Scientific), or polyethylenimine hydrochloride (PEI, Sigma Aldrich) in 1X OptiMEM (Gibco), according to the manufacturer’s instructions. For CRISPR/Cas9 KO generation, immunofluorescence, and live imaging experiments, each well was transfected with 0.5 µg of plasmid DNA. For co-immunoprecipitation experiments, the transfection was optimized for high efficiency and consistency by altering the plasmid concentration. All over-expression experiments were assessed 48 hours post transfection.

### Proximity dependent biotinylation-MS acquisition and analysis

All cell line generation and sample preparation were performed as previously described.^26^ Briefly, stable cell lines were made from Flp-In^TM^ T-REx^TM^293 cells (Thermo Fisher), which constitutively express the Tet repressor. To generate stable cell lines, cells were transfected with Lipofectamine 2000 (Thermo Fisher Scientific) and a combination of the appropriate Lis1 or Ndel1 BioID-fusion construct and pOG44, which expresses Flipase. After recovery, stables were selected with media supplemented with 50 µg/mL Hygromycin B. Colonies were isolated, expanded, and screened for expression of the fusion proteins by Western Blotting with an α-FLAG M2-HRP antibody (Sigma-Aldrich).

For BioID experiments, low passage cells were plated at 20% confluence in 15 cm dishes as 4 replicates, with each replicate consisting of 4 × 15 cm plates. After 24 hours, biotin was added to the media to a final concentration of 50 µM, and the cells were incubated for 16 hr. After removing the media, cells were removed from each plate by pipetting with ice-cold PBS and harvested via centrifugation at 1000 x g for 2 minutes. Cells were washed once more with ice cold PBS. To lyse, cells were resuspended in 8 mL RIPA buffer (50 mM Tris-HCl, pH 8.0; 150 mM NaCl, 1% (v/v) NP-40, 0.5% (w/v) sodium deoxycholate, 0.1% (w/v) SDS, 1 mM DTT, and protease inhibitors (cOmplete Protease Inhibitor Cocktail, Roche; Switzerland) by gentle rocking for 30 min at 4°C. The cell lysate was clarified via centrifugation at 30,000 rpm for 30 min in a Type 70 Ti rotor (Beckman Coulter) at 4°C. The clarified lysate was retrieved and incubated with 0.5 mL of streptavidin-conjugated beads (Dynabeads MyOne Streptavidin T1) and incubated overnight at 4°C with gentle rocking. Bead/lysate mixtures were collected on a magnetic stand and beads were then washed 3 times with 2 mL RIPA buffer. To elute immobilized proteins, the beads were boiled for 10 minutes at 100°C in 100 µL elution buffer (50 mM Tris, pH 6.8, 2% SDS (w/v), 20 mM DTT, 12.5 mM EDTA, 2 mM biotin). 90 µL of eluant was diluted to a final volume of 400 µL with 100 mM Tris-HCl, pH 8.5. 100 µL 100% trichloroacetic acid was then added and the solution was incubated overnight at 4°C. The precipitated sample was collected by centrifugation at maximum speed in a microcentrifuge for 30 minutes at 4°C. The pellet was washed twice with 500 µL ice cold 100% acetone. After removing the final acetone wash, the pellet was dried in a laminar flow cabinet for 30–60 minutes.

The samples were digested according to the FASP protocol using a 10 kDa molecular weight cutoff filter. In brief, the samples were mixed in the filter unit with 8 M urea, 100 mM M ammonium bicarbonate (AB) pH 8.0, and centrifuged at 14 000 *g* for 15 minites. The proteins were reduced with 10 mM DTT for 30 min at RT, centrifuged, and alkylated with 55 mM iodoacetamide for 30 min at RT in the dark. Following centrifugation, samples were washed 3 times with urea solution, and 3 times with 50 mM AB, pH 8.0. Protein digestion was carried out with sequencing grade modified Trypsin (Promega) at 1/50 protease/protein (wt/wt) at 37°C overnight. Peptides were recovered from the filter using 50mM AB. Samples were dried in Speed-Vac and desalted and concentrated on Thermo Scientific Pierce C18 Tip.

Samples were analyzed on a LTQ Orbitrap Velos mass spectrometer (Thermo Fisher Scientific) coupled to an Eksigent nanoLC-2D system through a nanoelectrospray LC −MS interface. A volume of 8 μl of sample was injected into a 10 μl loop using an autosampler. To desalt the sample, material was flushed out of the loop, loaded onto a trapping column (ZORBAX 300SB-C18, dimensions 5×0.3 mm×5 μm), and washed with 0.1% formic acid at a flow rate of 5 μL/min for 5 minutes. The analytical column was then switched on-line at 0.6 μl/min over an in-house made 100 μm i.d. × 200 mm fused silica capillary packed with 4 μm 80 Å Synergi Hydro C18 resin (Phenomex; Torrance, CA). After 10 min of sample loading, the flow rate was adjusted to 0.35 mL/min, and each sample was run on a 90-min linear gradient of 4–40% acetonitrile with 0.1% formic acid to separate the peptides. LC mobile phase solvents and sample dilutions used 0.1% formic acid in water (Buffer A) and 0.1% formic acid in acetonitrile (Buffer B) (Chromasolv LC–MS grade; Sigma-Aldrich, St. Louis, MO).

Data acquisition was performed using the instrument supplied Xcalibur™(version 2.1) software. The mass spectrometer was operated in the positive ion mode. Each survey scan of m/z 400–2000 was followed by collision assisted dissociation (CAD) MS/MS of twenty most intense precursor ions. Singly charged ions were excluded from CAD selection. Doubly charged and higher ions were included. Normalized collision energies were employed using helium as the collision gas.

MS/MS spectra were extracted from raw data files and converted into .mgf files using a Proteome Discoverer Software (ver. 2.1.0.62). These .mgf files were then independently searched against human database using an in-house Mascot server (Version 2.6, Matrix Science). Mass tolerances were +/- 10 ppm for MS peaks, and +/- 0.6 Da for MS/MS fragment ions. Trypsin specificity was used allowing for 1 missed cleavage. Methionine oxidation, protein amino-terminal acetylation, amino-terminal biotinylation, lysine biotinylation, and peptide amino-terminal pyroglutamic acid formation, were all allowed as variable modifications, while carbamidomethyl of Cysteine was set as a fixed modification. Scaffold (version 4.8, Proteome Software, Portland, OR, USA) was used to validate MS/MS based peptide and protein identifications. Peptide identifications were accepted if they could be established at greater than 95.0% probability as specified by the Peptide Prophet algorithm. Protein identifications were accepted if they could be established at greater than 99.0% probability and contained at least two identified unique peptides.

### CRISPR-Cas9 knockout cell line generation and validation

Guide sequences targeting exon 2 of CCSer2 were designed using CCTop Tool.^62^ Guides (Table 3) were inserted into px459 plasmid from Zhang lab using Bbs1 digestion as described.^63^ To generate KOs, U2OS WT cells were thawed and allowed to recover from freezing before plating 120,000 cells per well in a 6-well dish. The next day, cells were transfected with the px459 vector with or without CCSer2 guides, and one well was reserved for non-transfection control. Cells were allowed to grow for one day. Cells were then treated with 1 µg/mL puromycin to select for transfection. After complete cell death was achieved in the non-transfected well, the media was replaced with media lacking puromycin. After three days of clonal expansion, cells were collected for a surveyor assay and plated at a limiting dilution in 96-well dishes to select for individual cells. gDNA was collected from the cells transfected with px459 using the DNeasy kit (Qiagen), and PCR amplification followed by T7 endonuclease digestion confirmed that the guides cut efficiently. Only colonies from single cells were chosen for further amplification. Each clone was then screened for CCSer2-KO via western blot. To further confirm KO, cDNA was harvested from the isogenic KO cell lines and used for PCR amplification of the region surrounding the guide targeting site (Table 3). The generated PCR product was sequenced, and the sequencing results were analyzed using ICE Analysis (Synthego). To characterize the indels generated on each allele, PCR products generated for ICE analysis were TOPO (Thermo Fisher Scientific) cloned and sequenced as described.^64^ To ensure we captured every allele, we analyzed 30 TOPO clones.

### Co-Immunoprecipitation

60,000 U2OS cells were seeded per well of a 6-well dish and allowed to adhere overnight. Two wells of the 6-well dish were transfected with either vector control, CCSer2^WT^, or CCSer2^ΔCC^ using Lipofectamine LTX. The next day, the duplicate wells per sample were combined and re-plated onto a 10 cm TC-treated dish. Cells were allowed to grow for an additional day to reach confluency. Then, the cells were incubated for 10 minutes on ice with 1 mL of ice-cold Co-IP lysis buffer (30mM HEPES, (pH 7.4), 50mM KOAc, 0.1% Triton x100, 1mM DTT, 0.5mM Mg-ATP, 2 mM MgOAc, 1mM EGTA, 10% Glycerol, 1x Protease Inhibitor Cocktail (Roche)) and lysed with cell scrapers (Fisherbrand). Lysates were collected and centrifuged at max speed for 15 minutes at 4°C in a table-top microcentrifuge. 100 µL of supernatant was reserved for input and the remaining 900 µL was incubated with 30 µL anti-FLAG resin (Sigma-Aldrich A2220) for 4 hours with rocking at 4°C. After incubation, samples were spun down at 10,000 x g for 1 minute and the beads were washed four times with 1 mL cold Co-IP lysis buffer. After the final wash, beads were resuspended with 4X NuPage sample buffer (Fisher Scientific), 10% BME, and Co-IP lysis buffer to a volume of 100 µL. The samples were boiled for 5 minutes at 95°C and then used immediately for western blotting or frozen and stored at −20°C.

### Western blot

Samples from lysate or co-immunoprecipitation were loaded on a 4-12% BisTris NuPage gel (Thermo Fisher) and ran at 180 V for 60 minutes in 1x MOPS buffer (Thermo Fisher). The protein gels were then transferred onto methanol hydrated PVDF membranes (Amersham) for 3 hours at 300 mAmps at 4°C. Membranes were blocked with 5% nonfat milk in 1x TBST (Tris buffer saline with 0.05% Tween-20) with rocking for 30 minutes at room temperature. Membranes were then washed quickly with 1x TBST and primary antibodies diluted at various concentrations (Table 2) in 5% nonfat milk in 1x TBST were added to the membranes and incubated with rocking at 4°C overnight. Post incubation, membranes were washed extensively with 1x TBST before incubating with HRP-conjugated secondary antibodies for 1 hour at room temperature. Secondary antibodies were diluted at varying concentrations in 5% nonfat milk in 1x TBST (Table 2). Membranes were washed extensively with 1x TBST before visualization with either SuperSignal West Femto reagent (Thermo Fisher) or Clarity ECL reagent (BioRad) on the ChemiDoc imaging system (BioRad).

### Immunofluorescence

#1.5 22mm x 22mm coverslips (Corning) were washed in 100% ethanol and allowed to dry vertically in a 6-well dish. The coverslips were coated with either 1x Fibronectin human plasma (Sigma) or poly-D-lysine (20 µg/mL), incubated for 45 min at 37°C, washed with 1x PBS, and then allowed to dry completely or washed with 1x DMEM, respectively. Cells were plated at varying densities, depending on the downstream experimental requirement. Cells were fixed with either 100% ice cold methanol for 5 minutes on ice or 4% paraformaldehyde for 15 minutes at room temperature. After fixation, cells were washed with 3mL 1x PBS two times. Fixed cells were then blocked and permeabilized for 1 hour at room temperature while rocking with 1mL of blocking buffer (0.3% Triton X 100 and 5% normal goat serum (Cell Signaling Technology) in 1x PBS) and were washed quickly with 3mL 1x PBS. Primary antibodies were diluted in an antibody dilution buffer (0.1% Triton X100 and 0.5% BSA in 1x PBS) at varying concentrations (Table 2). The cells were incubated with the primary antibodies for 1 hour at room temperature. The coverslips were then washed extensively with 1x PBS for at least 15 minutes. Secondary antibodies and F-actin Phalloidin probes (Cayman Chemical) were diluted in antibody dilution buffer. The cells were incubated with the secondary antibodies and protected from light for 1 hour at room temperature. Stained coverslips were then washed quickly with 3mL 1x PBS. The cells were then treated with 1mL DAPI (1 µg/mL, Invitrogen) diluted in 1x PBS for 5 minutes, protected from light. The coverslips were then washed extensively with 1x PBS for at least 20 minutes. Washed coverslips were then mounted onto glass slides using one drop of ProLong Gold Antifade mountant (Fisher Scientific). Finally, the slides were cured for 24 hours in the dark and sealed with clear nail polish.

### Plasma membrane enrichment

To determine if CCSer2 displayed plasma membrane enrichment, WT U2OS cells transfected with CCSer2^WT^ using PEI were plated at a low confluency on fibronectin coated coverslips. The cells were then fixed with 4% PFA, stained for GFP and microtubules, and imaged with a spinning disk confocal microscope. Full volumes of cells were imaged by first identifying the top of each cell, then the bottom, then acquiring Z-slices at 0.2 µm steps. All analysis was performed in FIJI. A custom macro was written to partially automate the analysis. Using the GFP channel, the perimeter of the cell was outlined. An ROI band containing 2 µm on the interior of the perimeter and 0.25 µm outside the perimeter was generated and the intensity within this 2.25 µm perimeter band was measured in both the GFP and microtubule channels. To measure the cytoplasmic intensity, an ROI was generated that only included the area within the inner perimeter line. The ratio of the perimeter intensity to the cytoplasmic intensity was reported for both GFP-CCSer2 and microtubules. Only cells that were ∼98% in the field of view and not touching another cell on at least 75% of the perimeter were included in the analysis.

### +Tip analysis

To monitor plus-end localization of overexpressed CCSer2^WT^, CCSer2-SxNN^ALL^, and CCSer2-SxNN^1,4^ U2OS cells were transfected with 0.5 µg of plasmid using Lipofectamine LTX reagent (Thermo Fisher) as per manufacturers instruction. 24 hrs after transfections, cells were replated on into an 8-well live imaging dish (Thermo Fisher) coated with fibronectin. 24 hours after replating, cells were imaged via spinning disc confocal for 60 sec, 1 fps at 37°C and 5% CO_2_. All analysis was performed in FIJI. Maximum intensity projections of the first 15 frames of each movie were made and used for subsequent analysis. Whenever possible, the cell area was determined by manually thresholding each image. The perimeter was selected with the magic wand tool and area was measured. Next, all visible comets were counted manually in any cell where at least 3 comets were visible. Cells with excessive aggregation of CCSer2 constructs or no visible comets were excluded from the analysis. The number of comets counted for each cell was divided by area to get a measure of the density of comets per cell. Given the inherently subjective nature of this analysis, all analysis steps were performed blind.

### Cell shape analysis

To assess the elongated cell morphology we observed, WT and CCSer2 KO cells transfected with either vector, CCSer2^WT^, CCSer2^ΔCC^, or CCSer2-SxNN^ALL^ using Lipofectamine LTX and PLUS reagents (Invitrogen) and plated at low density onto PDL coated coverslips. Cells were fixed in 4% PFA, stained for actin with phalloidin-647, and imaged with a 20X objective spinning disk confocal microscope. Full volumes of cells were imaged by first identifying the top of each cell, then the bottom, then acquiring Z-slices at 0.9 µm steps. All analysis was performed in FIJI. A custom macro was written to partially automate the analysis. To begin, the actin channel was background subtracted and a gaussian blur was applied, then thresholding was used to identify individual cells. At this point, if two clearly individual cells were connected after thresholding by less than 25% of their cell periphery, then a black pencil drawn line was used to separate them. Only cells that were ∼98% in the field of view and not overlapping each other were included in the analysis. The magic wand tool was used to generate an ROI of each individual cell in the field of view. The circularity of each cell was then measured with the “shape descriptors” measurement in FIJI. The average circularity of the cells within one field of view was reported for each n.

### Projection behavior during nondirected cell migration

To assess the migratory behavior of individual cells during non-directed migration, ∼11,000 cells of either WT or CCSer2-KO cells were plated per well of an 8-well live imaging dish (Thermo Fisher) coated with fibronectin. 24 hours after plating, the cells were set up in an environmental chamber set at 37°C with 5% CO_2_ on a spinning disc confocal and allowed to acclimate for 30 minutes prior to imaging. Cells were imaged in DIC with a 20X objective for 24 hours at 1 frame per 3 minutes. All analysis was performed in the Nikon NIS-Elements software. From the 24-hour live imaging of WT and CCSer2-KO cells, the length and number of cellular projections made were quantified. For each cellular projection that was thinner than 10 µm and existed for at least 18 minutes, the maximum length was quantified by drawing a line segment from the base of the projection (at 10 µm thick) to the end of the projection. The percent of cells that formed projections within the 20X field of view was measured by dividing the number of cells that created at least one projection by the total number of cells in the field of view at the last frame of the movie.

### Particle Tracking – Nuclei tracking and directional persistence analysis

To monitor the directional persistence of individual cells during directed collective migration, ∼50,000 cells of either WT or CCSer2 KO cells were plated per well of a 2-well silicone insert (Ibidi) in an 8-well live imaging dish (LabTekII, Thermo Fisher) coated with fibronectin. 24 hours after plating, cells were incubated with 250nM SiR-DNA dye in 1x DMEM for 4 hours to stain the nuclei. The 2-well inserts were removed carefully with tweezers, the wells were gently washed twice with 1x PBS and then incubated with 1x DMEM with 100nM SiR-DNA dye. Cells were set up in an environmental chamber set at 37°C with 5% CO_2_ on a spinning disc confocal and allowed to acclimate for 30 minutes after removal of the insert. Four fields of view per well were imaged with a 10X objective for 24 hours at 1 frame per 3 minutes. All analysis was performed in FIJI. First, a 400 x 1064 µm rectangle ROI was drawn to encompass one half of the wound. All particle tracking was performed with the Trackmate plugin in FIJI.^65^ The nuclei were identified with the LoG detector. The particle diameter was set to 20 µm and the quality was set to 0.2 to select all nuclei within the field of view. The LAP Tracker was used to track nuclei movements. The max frame-frame linking distance was set to 20 µm with a quality penalty of 5. The max gap closing distance was set to 30 µm and the max frame gap was 2 with a quality penalty of 5. The cell tracks were then filtered by total distance traveled (above 70 µm), displacement (above 70 µm), and mean x position (+/- 52 µm, depending on the orientation of the wound). All track data was exported, then the directionality ratio (called confinement ratio in Trackmate) of each cell path was extracted and averaged per field of view. Directionality ratio is defined as the net displacement divided by the total distance the cell has migrated. The mean and max velocity of individual cells was also reported.

To assess the speed of collective cell migration during wound healing, the experimental set up was the same as above. Four fields of view per well were imaged in DIC with a 10X objective for 24 hours at 1 frame per 3 minutes. All analysis was performed in FIJI. The speed of both sides of the wound was calculated and then averaged to get the speed of wound closure per field of view. At time zero, a line was drawn down the center of the wound and at the leading edge of one half of the wound. The distance between these two lines was calculated. Once the cells crossed the threshold of the center line, the time was recorded. The distance traveled in microns over the time for the sheet of cells to close the wound was reported as the speed of wound closure. This analysis was repeated on the other half of the wound and the speed was averaged for both collectively migrating sheets of the wound.

### Anaphase onset analysis

To monitor the mitotic progression in WT and CCSer2 KO cells, ∼11,000 cells of either WT or CCSer2 KO cells were plated per well of an 8-well live imaging dish (Thermo Fisher) coated with fibronectin. 24 hours after plating, the cells were set up in an environmental chamber set at 37°C with 5% CO_2_ on a spinning disc confocal and allowed to acclimate for 30 minutes prior to imaging. Cells were imaged in DIC with a 20X objective for 24 hours at 1 frame per 3 minutes. All analysis was performed in FIJI. From the 24-hour live imaging of WT and CCSer2-KO cells, each cell that completed cell division within the time constraints of the movie was analyzed. Using a mitotic phase guide created from WT U2OS cells before starting the analysis, the time from the beginning of prophase to the first separation of the chromosomes during anaphase was recorded.

### Wound healing and polarization assay

Cell polarity assays were performed largely as described.^41,66^ Briefly, 70,000 cells were seeded onto fibronectin coated coverslips and were allowed to acclimate overnight at 37°C with 5% CO2. Cells at confluency were then starved for 24 hours in 3mL DMEM starvation media (1x DMEM with 1% Penicillin/ Streptomycin and no FBS) to synchronize cells. Starved cells were then washed with 3mL 1X PBS (Gibco) and scratched with a p200 tip in a cross pattern. 3mL of migration media (1x DMEM with 1% FBS, and 1% Penicillin/Streptomycin) was then added to each well. The wounded cells were allowed to migrate for 0 hours, 30 minutes, or 4 hours at 37°C with 5% CO2 before fixation with either methanol or 4% PFA. After fixation, cells were washed with 3mL 1X PBS three times and prepared for immunofluorescence as previously described. For live imaging experiments of wound closure, cells were instead plated on fibronectin-coated 8-well dishes (Lab-Tek). Wounds were either generated as described above or with a 2-well silicone insert (ibidi). Wound closure was visualized for 16-24 hours at 37°C and 5% CO_2_.

### Ndel1 and dynein cortical enrichment during directed migration analysis

To assess Ndel1 and dynein localization during migration, confluent U2OS cells on fibronectin coated coverslips were fixed in ice cold 100% methanol for 5 minutes and stained with α-tubulin and either α-Ndel1 or α-dynein heavy chain antibodies, respectively. Cells were imaged with a 60X objective spinning disk confocal microscope. Full volumes of cells were imaged by first identifying the top of each cell, then the bottom, then acquiring Z-slices at 0.2 µm steps. All analysis was done in FIJI. A custom macro was written to partially automate the analysis. First, the user provides the correct orientation of the image then using the polyline tool draws a line at the cell border in the dynein channel. The line contains many points that are evenly spaced to ensure proper measurement of the cytoplasmic dynein levels in the next step. After the line is drawn, the macro will expand the line by 5 pixels (∼1 µm) to measure cortex intensity and generate 15 circle ROIs with a 15 pixel diameter (2.75 µm), 35 pixels (6.416 µm) behind the previously defined cell border in the interior of the cell to measure cytoplasmic dynein intensity. The user is then prompted to delete any overlapping circles. The circles are then combined into one ROI and the area and mean intensity at the cortex and in the cytoplasm are measured in both the dynein and tubulin channels.

### Particle Tracking – Integrin vesicle trafficking analysis

To monitor integrin containing vesicle movements, U2OS WT and CCSer2 KO cells were transfected with mCherry-α5-integrin12 in a 6-well dish using Lipofectamine LTX. 24 hours after transfection, ∼65,000 cells were replated per well of an 8-well live imaging dish (Thermo Fisher) coated with fibronectin. 24 hours after replating, confluent cells were scratched with a p200 pipette tip, washed twice with 1x PBS, and once with warmed Live Cell Imaging Solution (Invitrogen). Cells were imaged an hour after wounding via spinning disc confocal with a 60X objective for 2 minutes at 2 fps in an environmental chamber set at 37°C with 5% CO_2_. All analysis was performed in FIJI. A custom macro (provided by John Salogiannis) was written to semi-automate the analysis prior to using the Trackmate plugin. Each movie was oriented with the cells migrating towards the bottom of the image. In this way, vesicles moving downward are trafficked in the anterograde direction and vesicles moving upward are trafficked in the retrograde direction. The background was subtracted with a 50 pixel rolling ball radius, the contrast was increased to a saturation point of 0.35, and the movie was converted to 32-bit. Next, using the line tool, a 5 µm line was drawn through an individual vesicle on the cell periphery, so that the vesicle is centered on the line. From the plot profile of the line, we fitted a gaussian curve and identified the d parameter. A gaussian blur in scaled units was applied to the movie using the measured d parameter as the radius. Then, the Mexican Hat Filter plugin was applied to the movie with a radius of 2. To limit the analysis to one cell, an ROI was drawn to isolate the space between the nucleus and the leading edge of a single cell. All tracking was performed with the Trackmate plugin in FIJI.^65^ The integrin vesicles were identified with the LoG detector. The particle diameter was set to 0.6 and the quality was set between 15 and 30 to select all vesicles within the ROI. The LAP Tracker was used to track integrin vesicles. The max frame-frame linking distance was set to 3 µm with a quality penalty of 5. The max gap closing distance was set to 2 µm and the max frame gap was 1 with a quality penalty of 5. The tracks were then filtered by linearity of forward progression (above 0.1), displacement (above 1 µm), and duration (above 5 seconds). All track data was exported as well as the MotilityLab spreadsheet data which included the XY positions of each spot within the tracks. A custom macro was written in excel to calculate the sign of the y-value track displacement, by subtracting the final y-position from the starting y-position of each track. Vesicles tracks were sorted by sign with a negative y displacement having traveled in the retrograde direction and a positive y displacement having traveled in the anterograde direction.

### Cargo distribution and EGF localization analysis

To assess the radial localization of different organelle cargoes inside the cell, WT and CCSer2 KO cells were fixed in 4% PFA, stained for the golgi apparatus, mitochondria, ER, early endosomes, lysosomes, and actin with α-GM130, TOM20, PDI, EEA1, LAMP1 antibodies and a phalloidin-647 probe, respectively. Cells were imaged with a 60X objective on a spinning disk confocal microscope. Full volumes of cells were imaged by first identifying the top of each cell, then the bottom, then acquiring z-slices at 0.2 µm steps. All analysis was done in FIJI. A custom macro was written to automate the analysis. First, a single cell was cropped using the rectangle tool. The channels were then split and the nucleus was identified and selected using automated thresholding and particle analysis tools. Next, the cell periphery was traced in the actin channel and saved as an ROI. From the centroid of the nucleus, lines were drawn to connect to the cell perimeter radially in intervals of 1 degree. The length of each line was measured and only lines longer than half of the maximum line were kept for analysis. The intensity across each line in the selected cargo channel was measured and binned into 20 sections. Finally, each bin value was averaged across all the line measurements so that a fractional intensity distribution profile was created for one cell. These distribution profiles were then combined across cells and bioreplicates to generate a final normalized plot.

To assess the radial localization of EGF-555 puncta inside the cell, WT and CCSer2 KO cells were treated with EGF-555 ligand for 5 minutes and chased for either 0 or 30 minutes. Cells were washed extensively with 1x PBS and fixed in 4% PFA, stained for actin with a Phalloidin-647 probe. Cells were imaged with a 60X objective on a spinning disk confocal microscope. Full volumes of cells were imaged by first identifying the top of each cell, then the bottom, then acquiring z-slices at 0.2 µm steps. All analysis was done in FIJI. The custom macro written for the cargo distribution analysis was used for the EGF-555 localization. The distribution profiles generated were compiled across cells and bioreplicates to generate a final normalized plot of EGF-555 distribution 5 minutes or 30 minutes after treatment.

### Early endosomes mean intensity analysis

To quantify the total mean intensity of early endosomes per cell, WT and CCSer2 KO cells were fixed in 4% PFA, stained for early endosomes and actin with an α-EEA1 antibody and a phalloidin-647 probe, respectively. Cells were imaged with a 60X objective on a spinning disk confocal microscope. Full volumes of cells were imaged by first identifying the top of each cell, then the bottom, then acquiring z-slices at 0.2 µm steps. All analysis was done in FIJI. A custom macro was written to automate the analysis. First, using the freehand tool, the cell periphery of one cell is traced in the actin channel and saved as an ROI. The cell perimeter ROI was then applied to the early endosome channel and the area and mean intensity were measured. The mean intensity of the early endosomes contained within a single cell was reported for each n.

### Zebrafish husbandry and ccser2 knockout generation

All zebrafish (*Danio rerio*) work was done in accordance with the University of Wisconsin-Madison IACUC guidelines. Zebrafish lines used include: AB, TgBAC(*neurod:egfp*)^nl1^, *ccser2a ^uwd^*^10^ *and ccser2b ^uwd^*^11^. Adult zebrafish were kept at 28°C and spawned according to established protocols^67^. Embryos and larval zebrafish were kept at 28°C in embryo media. Developmental staging was done according to established methods.^68^ Experiments were done at 4 dpf at which point sex is not determined.

To generate the stable *ccser2a* and *ccser2b* knockout zebrafish lines, two guide RNAs targeted to exon two of each gene (Table 3) were injected into zebrafish zygotes with 500 ng Cas9 protein (Integrated DNA Technologies). F0 injected animals were raised to adulthood and crossed to AB. A subset of the resulting larvae were genotyped with *ccser2a* or *ccser2b* specific primers to confirm indel creation (Table 3). The rest of the F1 larvae were raised to adulthood and then screened for frameshift deletions using PCR-based genotyping and sequencing. This identified *ccser2a ^uwd^*^10^ which has a 156 bp deletion and *ccser2b ^uwd^*^11^ which has a 446 bp deletion. Both alleles insert premature stop codons in the respective genes. Resulting heterozygous F1 animals were crossed to generate *ccser2a ^uwd^*^10^*^/+^*; *ccser2b ^uwd^*^11^*^/+^* double heterozygous F2s for analysis.

### G0 crispant and morphant analysis

For G0 crispant analysis, all four guide RNAs described above targeting *ccser2a* and *ccser2b* were coinjected with 500 ng Cas9 protein (Integrated DNA Technologies) into TgBAC(*neurod:egfp*)^nl1^ transgenics. Animals were raised to 4 dpf prior to immunofluorescence which was done according to established protocols.^67^ Briefly, animals were incubated in 4% PFA with 0.1% triton overnight at 4°C. Following fixation, larvae were washed in 1X PBS/0.1% triton briefly then washed in RNAse/DNAse-free water overnight at room temperature. Larvae were then incubated in standard blocking solution (5% goat serum, 0.2% triton, 1X PBS, 1% DMSO, 0.02% sodium azide, 0.2% bovine serum albumin) for 1-4 hours then incubated in primary antibody (Aves; GFP-1020) in blocking solution overnight at 4°. Following incubation in primary antibody, larvae were washed in 1X PBS/0.1% triton, then incubated in Alexa Fluor secondary antibodies (Invitrogen; 1:1000) and DAPI overnight at 4°. Larvae were washed again in 1X PBS/0.1% triton then sunk in 60% glycerol. Immunolabeled larvae were then imaged on an Olympus FV3000, 10x/NA0.4 or 60X/NA1.4 oil immersion objective.

For *ccser2a;ccser2b* morphant analysis, 1 ng of each morpholino targeting the start site of *ccser2a* and *ccser2b* (Table 3) were co-injected into TgBAC(*neurod:egfp*)^nl1^ transgenic zygotes. Animals were raised to 30 hpf or 4 dpf and processed for immunofluorescence as described above. Immunolabeled larvae were then imaged on an Olympus FV3000, 10X/NA0.4 or 60x/NA1.4 oil immersion objective.

Image analysis was performed in ImageJ. Neuromast number was manually counted in projected stacks from 10X images. pLLp area and number of mitotic and apoptotic cells were measured from 60X images. For pLLp area, the primordium was manually outlined and area measured using built in plugins in ImageJ. Number of mitotic and apoptotic nuclei were manually counted in the *z*-stacks. For pLL nerve to body length ratios, the length of the lateral line nerve and the length of the body from the ear to the tail were measured using built in plugins in ImageJ.

### In situ hybridization and RT-PCR analysis

cDNA was synthesized from total RNA extracted from 2 cell stage zebrafish embryos (for RT-PCR) or from 4 dpf larvae (for in situ hybridization). Total RNA was extracted using Trizol reagent according to the manufacturer’s instructions (Invitrogen). cDNA synthesis was done using a Superscript IV reverse transcription kit according to the manufacturer’s instructions (Invitrogen).

In situ hybridization and probe synthesis were done according to established protocols.^69,70^ DIG-labeled PCR-based probe templates for *ccser2a* and *ccser2b* (Table 3) were synthesized from 4 dpf cDNA generated as described above.

For RT-PCR analysis of *ccser2a* and *ccser2b* mRNA levels, cDNA was synthesized from 2 cell stage zebrafish embryos. This developmental stage is before the onset of zygotic transcription and therefore represents maternally deposited mRNA (Table 3).

### Protein expression and purification

The His-ZZ-TEV-Halo-CCSer2^658–850^ and His-Strep-sfGFP-CCSer2^650–850^ were expressed from the pET28 vector backbone in BL21 (DE3) *E.coli* cells after induction with 0.5 mM Isopropyl ß-D-1-thiogalactopyranoside (IPTG) for 18 hours at 16 °C and 200 rpm. The induced cells were harvested by centrifugation (6000 x g, 20 minutes, 4 °C). The pellets were resuspended in lysis buffer (30 mM HEPES [pH 7.4], 50 mM potassium acetate, 2 mM magnesium acetate, 1 mM EGTA, 1 mM DTT, 0.5 mM Pefabloc, 10% [v/v] glycerol) supplemented with 1× cOmplete EDTA-free protease inhibitor cocktail tablets (Roche). The resuspended cells were incubated on ice for 30 min with 1 mg/mL egg lysozyme and sonicated (50% amplitude, pulse on 5 s, pulse off 25 s). The lysate was clarified by centrifugation (66,000 x g, 30 minutes, 4 °C) in a Type 70Ti rotor (Beckman). For His-ZZ-TEV-Halo-CCSer2^650–850^, the supernatant was incubated with 2 mL of IgG Sepharose 6 Fast Flow beads (Cytiva) equilibrated in lysis buffer and incubated for 2 hours with rotation at 4 °C. The beads were collected by centrifugation (1000 x g, 2 minutes, 4 °C) and resuspended with 2 mL lysis buffer before being transferred to a glass gravity column. Beads were then washed with 50 mL low salt wash buffer (30 mM HEPES [pH 7.4], 200 mM potassium acetate, 2 mM magnesium acetate, 1 mM EGTA, 1 mM DTT, 0.5 mM Pefabloc, 10% [v/v] glycerol), 100 mL high salt wash buffer (30 mM HEPES [pH 7.4], 1050 mM potassium acetate, 2 mM magnesium acetate, 1 mM EGTA, 1 mM DTT, 0.5 mM Pefabloc, 10% [v/v] glycerol), 200 mL low salt wash buffer and 100 mL tobacco etch virus (TEV) buffer (50 mM Tris–HCl [pH 8.0], 150 mM potassium acetate, 2 mM magnesium acetate, 1 mM EGTA, 1 mM DTT and 10% [v/v] glycerol). The ZZ-TEV tag was cleaved by incubating the beads with TEV protease at a final concentration of 0.2 mg/mL overnight. Cleaved Halo-CCSer2^650–850^ was concentrated to 1 mL with a 30K molecular weight cut off (MWCO) concentrator (EMD Millipore) and diluted with 1 mL buffer A (30 mM HEPES [pH 7.4], 50 mM potassium acetate, 2 mM magnesium acetate, 1 mM EGTA, 1 mM DTT, 10% [v/v] glycerol). The 2 mL protein sample was loaded into a MonoQ 5/50 GL column (Cytiva) at 1 mL/min. The column was prewashed with 10 CVs of buffer A, 10 CVs of buffer B (30 mM HEPES [pH 7.4], 200 mM potassium acetate, 2 mM magnesium acetate, 1 mM EGTA, 1 mM DTT, 10% [v/v] glycerol) and again with 10 CVs of buffer A. To elute, a linear gradient was run over 26 CVs from 0 to 100% buffer B. The peak fractions were collected and concentrated to 500 μL with a 30K MWCO concentrator (EMD Millipore). The 500 µL protein sample was further subjected to size exclusion chromatography (SEC) on a Superdex200 10/300 column (Cytiva) with GF150 buffer (25 mM HEPES [pH 7.4], 150 mM KCl, 1 mM MgCl2, 1 mM DTT) as the mobile phase at 0.75 mL/min. Peak fractions were collected, supplemented with glycerol to a final concentration of 10%, concentrated to 0.2–1 mg/mL with a 30K MWCO concentrator (EMD Millipore), frozen in liquid nitrogen, and stored at –80°C. For fluorescent labeling of purified Halo-CCSer2^650–850^, the Halo-CCSer2^650–850^ was mixed with a 10-fold excess of Halo-TMR (Promega) for 10 min at room temperature. Unconjugated dye was removed by passing the protein through Micro Bio-spin P-6 column (Bio-rad) equilibrated in GF150 buffer supplemented with 10% glycerol. Small volume aliquots of the labeled protein were flash-frozen in liquid nitrogen and stored at −80°C.

For the His-Strep-sfGFP-CCSer2^650–850^, the purification was similar to ZZ-TEV-Halo-CCSer2^650–850^. Briefly, 2 mL of HisPurTM Ni-NTA resin was used. After incubation overnight, the beads were washed with 50 mL low salt wash buffer supplemented with 25mM imidazole, 100 mL high salt wash buffer supplemented with 25mM imidazole, and 200 mL low salt wash buffer supplemented with 25mM imidazole. The proteins were then eluted with 10 mL elution buffer (30 mM HEPES [pH 7.4], 50 mM potassium acetate, 2 mM magnesium acetate, 300mM imidazole, 1 mM EGTA, 1 mM DTT, 10% [v/v] glycerol) for 15min at 4°C. The eluted proteins were concentrated to 1 mL with a 30K MWCO concentrator (EMD Millipore) and diluted with 1 mL buffer A. The 2mL protein sample was loaded into a MonoQ 5/50 GL column (Cytiva) at 1 mL/min. The column was prewashed with 10 column volumes (CVs) of buffer A, 10 CVs of buffer B and again with 10 CVs of buffer A. To elute, a linear gradient was run over 26 CVs from 0 to 100% buffer B. The peak fractions were collected and concentrated to 500 µL with a 30K MWCO concentrator (EMD Millipore). The 500 µL protein sample was further subjected to SEC on a Superdex200 10/300 column (Cytiva) with GF150 buffer as the mobile phase at 0.75 mL/min. Peak fractions were collected, supplemented with glycerol to a final concentration of 10%, concentrated to 0.2–1 mg/mL with a 30K MWCO concentrator (EMD Millipore), frozen in liquid nitrogen, and stored at –80°C.

Human dynein, Lis1, Ndel1 and NT-Ndel1 constructs were expressed in Sf9 cells as described.^6,21,71^ Briefly, pACEBac1 plasmid containing the human dynein genes, pFastBac plasmid containing Lis1, and pKL plasmid containing Ndel1 and tagged Lis1 constructs were transformed into DH10EmBacY chemically competent cells with heat shock at 42 °C for 15 seconds followed by incubation at 37 °C and shaking at 220 rpm for 6 hours in S.O.C. media (Thermo Fisher Scientific). The cells were plated on LB–agar plates containing kanamycin (50 μg/mL), gentamicin (7 μg/mL), tetracycline (10 μg/mL), Bluo-Gal (100 μg/mL), and IPTG (40 μg/mL). Cells that contained the plasmid of interest were identified with blue/white selection after 48 to 72 h. For dynein, white colonies were tested for the presence of all six dynein genes with PCR. Colonies were grown overnight in LB medium containing kanamycin (50 μg/mL), gentamicin (7 μg/mL), and tetracycline (10 μg/mL) at 37 °C and agitation at 220 rpm. Bacmid DNA was extracted from overnight cultures using isopropanol precipitation as described.^72^ About 1 × 106 Sf9 cells in 2 mL of media in a 6-well dish were transfected with up to 2 μg of fresh bacmid DNA using FuGene HD transfection reagent (Promega) at a ratio of 3:1 (Fugene reagent:DNA) according to the manufacturer’s directions. Cells were incubated at 27 °C for 3 days without agitation in a humid incubator. Next, the supernatant containing the virus (V0) was harvested by centrifugation (1000 g, 5 minutes, 4 °C). About 1 mL of the V0 virus was used to transfect 50 mL of Sf9 cells at 10^6^ cells/mL to generate the next passage of virus (V1). Cells were incubated at 27 °C for 3 days with shaking at 105 rpm. Supernatant containing V1 virus was collected by centrifugation (1000 g, 5 minutes, 4 °C). All V1 were protected from light and stored at 4 °C until further use. To express protein, 4 mL of V1 virus were used to transfect 400 mL of Sf9 cells at a density of 1 × 106 cells/mL. Cells were incubated at 27 °C for 3 days with shaking at 105 rpm and collected by centrifugation (3500 g, 10 minutes, 4 °C). The pellet was washed with 10 mL of ice-cold PBS and collected again via centrifugation before being flash frozen in liquid nitrogen and stored at −80 °C until needed for protein purification.

All steps for protein purification were performed at 4 °C unless indicated otherwise. For dynein preparation, Sf9 cell pellets were thawed on ice and resuspended in 40 mL of dynein-lysis buffer (50 mM HEPES [pH 7.4], 100 mM sodium chloride, 1 mM DTT, 0.1 mM Mg–ATP, 0.5 mM Pefabloc, 10% [v/v] glycerol) supplemented with one cOmplete EDTA-free protease inhibitor cocktail tablet (Roche) per 50 mL. Cells were lysed with a Dounce homogenizer (ten strokes with a loose plunger followed by 15 strokes with a tight plunger). The lysate was clarified by centrifugation (183,960 x g, 88 minutes, 4 °C) in a Type 70Ti rotor (Beckman). The supernatant was mixed with 2 mL of IgG Sepharose 6 Fast Flow beads (Cytiva) equilibrated in Dynein-lysis buffer and incubated for 4 hours with rotation along the long axis of the tube. The beads were transferred to a glass gravity column, washed with at least 200 mL of dynein-lysis buffer and 300 mL of Dynein-TEV buffer (50 mM Tris–HCl [pH 8.0], 250 mM potassium acetate, 2 mM magnesium acetate, 1 mM EGTA, 1 mM DTT, 0.1 mM Mg–ATP, and 10% [v/v] glycerol). For fluorescent labeling of SNAP tag, dynein-bound beads were mixed with 5 μM SNAP-Cell-TMR or SNAP-AlexaFluor-647 (New England Biolabs) for 10 minutes at room temperature.

Unconjugated dye was removed by washing with 300 mL Dynein-TEV buffer at 4 °C. The beads were resuspended in 15 mL of TEV buffer supplemented with 0.5 mM Pefabloc and 0.2 mg/mL TEV protease and incubated overnight with rotation along the long axis of the tube. Cleaved proteins in the supernatant were concentrated with a 100K (MWCO) concentrator (EMD Millipore) to 500 μL and purified via SEC on a TSKgel G4000SWXL column (TOSOH Bioscience) with GF150 buffer supplemented with 0.1 mM Mg–ATP as the mobile phase at 0.75 mL/min. Peak fractions were collected, buffer exchanged into a GF150 buffer supplemented with 0.1 mM Mg–ATP and 10% glycerol, and concentrated to 0.1 to 0.5 mg/mL using a 100K MWCO concentrator. Small-volume aliquots were flash frozen in liquid nitrogen and stored at −80 °C.

Lysis and clarification steps for Lis1 and Ndel1 constructs were similar to dynein except Lis1-lysis buffer (30 mM HEPES [pH 7.4], 50 mM potassium acetate, 2 mM magnesium acetate, 1 mM EGTA, 300 mM potassium chloride, 1 mM DTT, 0.5 mM Pefabloc, 10% [v/v] glycerol) supplemented with one cOmplete EDTA-free protease inhibitor cocktail tablet per 50 mL was used in place of dynein-lysis buffer. The clarified supernatant was mixed with 2 mL of IgG Sepharose 6 Fast Flow beads (Cytiva) and incubated for 2 to 3 hours with rotation along the long axis of the tube. The beads were transferred to a gravity column, washed with at least 20 mL of Lis1-lysis buffer, 200 mL of Lis1-TEV buffer (10 mM Tris–HCl [pH 8.0], 2 mM magnesium acetate, 150 mM potassium acetate, 1 mM EGTA, 1 mM DTT, 10% [v/v] glycerol) supplemented with 100 mM potassium acetate and 0.5 mM Pefabloc, and 100 mL of Lis1-TEV buffer. For fluorescent labeling of Lis1, the Lis1-bound beads were mixed with SNAP-AlexaFluor 647 (Promega) for 10 minutes at room temperature after the lysis buffer wash. TEV protease was added to the beads at a final concentration of 0.2 mg/mL, and the beads were incubated overnight with rotation along the long axis of the tube. Cleaved Lis1 in the supernatant was collected and concentrated to 500 µL with a 30K MWCO concentrator (EMD Millipore). Concentrated Lis1 was then purified via SEC on a Superose 6 Increase 10/300 GL column (Cytiva) with GF150 buffer supplemented with 10% glycerol as the mobile phase at 1 ml/min. Peak fractions were collected, concentrated to 0.2–1 mg/mL with a 30K MWCO concentrator (EMD Millipore), frozen in liquid nitrogen, and stored at –80°C.

For the GST-Ndel1 constructs, 2 mL of HisPurTM Ni-NTA resin was used. After incubation overnight, the beads were washed with 50 mL low salt wash buffer supplemented with 25mM imidazole, 100 mL high salt wash buffer supplemented with 25mM imidazole, and 200 mL low salt wash buffer supplemented with 25 mM imidazole. The proteins were then eluted with 10 mL elution buffer for 15 minutes at 4°C. The eluted proteins were collected and concentrated to 500 µL with a 30K MWCO concentrator (EMD Millipore). Concentrated Ndel1 was then purified via SEC on a Superdex200 10/300 column (Cytiva) with GF150 buffer supplemented with 10% glycerol as the mobile phase at 1 mL/min. Peak fractions were collected, concentrated to 0.2–1 mg/mL with a 30K MWCO concentrator (EMD Millipore), frozen in liquid nitrogen, and stored at –80°C.

Dynactin was purified from human embryonic kidney 293T cell lines stably expressing p62-Halo-3xFLAG as described.^26^ Briefly, frozen pellets collected from 160 x 15 cm plates were resuspended in 80 mL of dynactin-lysis buffer (30 mM HEPES [pH 7.4], 50 mM potassium acetate, 2 mM magnesium acetate, 1 mM EGTA, 1 mM DTT, 10% [v/v] glycerol) supplemented with 0.5 mM Mg-ATP, 0.2% Triton X-100, and 1× cOmplete EDTA-free protease inhibitor cocktail tablets) and rotated along the long axis of the tube for at least 15 minutes. The lysate was clarified via centrifugation (66,000 x g, 30 minutes, 4 °C) in a Type 70 Ti rotor (Beckman). The supernatant was mixed with 1.5 mL of α-FLAG M2 affinity gel (Sigma–Aldrich) and incubated overnight with rotation along the long axis of the tube. The beads were transferred to a glass gravity column, washed with at least 50 ml of wash buffer (dynactin-lysis buffer supplemented with 0.1 mM Mg–ATP, 0.5 mM Pefabloc, and 0.02% Triton X-100), 100 mL of wash buffer supplemented with 250 mM potassium acetate, and then washed again with 100 mL of wash buffer. About 1 mL of FLAG elution buffer (wash buffer with 2 mg/mL of 3xFLAG peptide) was used to elute dynactin, which was then filtered via centrifuging through an Ultrafree-MC VV filter (EMD Millipore) in a tabletop centrifuge according to the manufacturer’s instructions. The filtered dynactin was then diluted to 2 mL in dynactin buffer A (50 mM Tris–HCl [pH 8.0], 2 mM magnesium acetate, 1 mM EGTA, and 1 mM DTT) and loaded onto a MonoQ 5/50 GL column (Cytiva) at 1 mL/min. The column was prewashed with 10 column volumes (CVs) of dynactin buffer A, 10 CVs of dynactin buffer B (50 mM Tris–HCl [pH 8.0], 2 mM magnesium acetate, 1 mM EGTA, 1 mM DTT, 1 M potassium acetate) and then equilibrated with 10 CVs of dynactin buffer A. A linear gradient was run over 26 CVs from 35 to 100% buffer B. Pure dynactin eluted between 75 and 80% buffer B. Peak fractions were collected, pooled, buffer exchanged into a GF150 buffer supplemented with 10% glycerol, concentrated to 0.02 to 0.1 mg/mL using a 100K MWCO concentrator, aliquoted into small volumes, and then flash frozen in liquid nitrogen.

BicD2 (amino acids 25 – 398) containing an amino-terminal HaloTag were expressed in BL-21[DE3] cells till they reached an optical density of 0.4 to 0.6 at 600 nm. At that point protein expression was induced with 0.1 mM IPTG for 16 hours at 18°C. Frozen cell pellets from a 1.5 L culture were resuspended in 40 mL of activating-adaptor-lysis buffer (30 mM HEPES [pH 7.4], 50 mM potassium acetate, 2 mM magnesium acetate, 1 mM EGTA, 1 mM DTT, and 0.5 mM Pefabloc, 10% [v/v] glycerol) supplemented with 1× cOmplete EDTA-free protease inhibitor cocktail tablets and 1 mg/mL lysozyme. The resuspension was incubated on ice for 30 minutes and lysed by sonication. The lysate was clarified by centrifuging at 66,000g for 30 minutes in Type 70 Ti rotor. The clarified supernatant was incubated with 2 mL of IgG Sepharose 6 Fast Flow beads (Cytiva) for 2 hours on a roller. The beads were transferred into a gravity-flow column, washed with 100 mL of activating-adaptor-lysis buffer supplemented with 150 mM potassium acetate and 50 mL of cleavage buffer (50 mM Tris– HCl [pH 8.0], 150 mM potassium acetate, 2 mM magnesium acetate, 1 mM EGTA, 1 mM DTT, 0.5 mM Pefabloc, and 10% [v/v] glycerol). The beads were then resuspended and incubated in 15 mL of cleavage buffer supplemented with 0.2 mg/mL TEV protease overnight with rotation along the long axis of the tube. The supernatant containing cleaved proteins was concentrated using a 30K MWCO concentrator to 1 mL, filtered by centrifuging with Ultrafree-MC VV filter (EMD Millipore) in a tabletop centrifuge, diluted to 2 mL in buffer A and injected into a MonoQ 5/50 GL column (Cytiva) at 1 mL/min. The column was prewashed with 10 CVs of buffer A, 10 CVs of buffer B and again with 10 CVs of buffer A. To elute, a linear gradient was run over 26 CVs from 0 to 100% buffer B. The peak fractions containing Halo-tagged activating adaptors were collected and concentrated to using a 30K MWCO concentrator to 0.2 mL, diluted to 0.5 mL in GF150 buffer, and further purified using size-exclusion chromatography on a Superose 6 Increase 10/300 GL column (Cytiva) with GF150 buffer at 0.5 mL/min. The peak fractions were collected, buffer-exchanged into a GF150 buffer supplemented with 10% glycerol, concentrated to 0.2 to 1 mg/mL using a 30K MWCO concentrator, and flash frozen in liquid nitrogen.

### Single Molecule TIRF microscopy data acquisition, analysis, and statistical tests

Single-molecule imaging was performed with an inverted microscope (Nikon, Ti2-E Eclipse) with a 100x 1.49 N.A. oil immersion objective (Nikon, Apo). The microscope was equipped with a LUNF-XL laser launch (Nikon), with 405 nm, 488 nm, 561 nm, and 640 nm laser lines. The excitation path was filtered using an appropriate quad bandpass filter cube (Chroma). The emission path was filtered through appropriate emission filters (Chroma) located in a high-speed filter wheel (Finger Lakes Instrumentation). Emitted signals were detected on an electron-multiplying CCD camera (Andor Technology, iXon Ultra 897). Image acquisition was controlled by NIS Elements Advanced Research software (Nikon).

Single-molecule motility and microtubule binding assays were performed in flow chambers assembled as described previously.^71^ No. 1-1/2 coverslips (Corning) were functionalized with biotin-PEG by sonication at 40°C with 100% EtOH for 10 minutes, 200mM KOH for 20 minutes, and again with 100% EtOH for 10 minutes. Coverslips were rinsed with water 3 times in between sonication steps. After the last EtOH wash the coverslips were incubated with methanol containing 5% acetic acid and 1% (3-Aminopropyl)triethoxysilane (Millipore-Sigma) overnight in a vacuum desiccation chamber in the dark. The next day the coverslips were washed with water 5 times and once with 100% EtOH before drying. Coverslips were incubated with 8.4mg/mL NaHCO3, 270 mg/mL mPEG-Succinimidyl Valerate, MW 2,000 (Laysan Bio) and 35 mg/mL Biotin-PEG-SVA, MW 5,000 (Laysan Bio) in a humid container for 3 hours. Slides were washed with water, dried, and stored at −20°C or in a vacuum desiccator.

Taxol-stabilized microtubules with ∼10% biotin-tubulin and ∼10% Alexa488 labelled fluorescent-tubulin were prepared as described.^73^ Flow chambers were assembled with taxol-stabilized microtubules by incubating 1 mg/mL streptavidin in assay buffer (30 mM HEPES [pH 7.4], 2 mM magnesium acetate, 1 mM EGTA, 10% glycerol, 1 mM DTT) for 3 minutes, washing twice with assay buffer supplemented with taxol, incubating a fresh dilution of taxol-stabilized microtubules in assay buffer for 3 minutes, and washing twice with assay buffer supplemented with 1 mg/mL casein and 20 µM Taxol.

Dynein was incubated in the presence of CCSer2^650–850^, Ndel1, Lis1, and/or GF150 buffer (to buffer match for experiments without CCSer2, Ndel1, or Lis1) for 10 minutes on ice. These complexes were then incubated an additional 10 minutes on ice with dynactin and BicD2 at a ratio of 1:2:10 for dynein:dynactin:BicD2. They were then flowed into the flow chamber assembled with taxol-stabilized microtubules. The final imaging buffer contained the assay buffer supplemented with 20 µM Taxol, 1 mg/mL casein, 71.5 mM β-mercaptoethanol, 0.05 mg/mL glucose catalase, 1.2 mg/mL glucose oxidase, 0.4% glucose, and 2.5 mM Mg-ATP. The final concentration of dynein in the flow chamber was 0.5-1 nM. The final imaging buffer contained 37.5 mM KCl.

For single-molecule motility assays, microtubules were imaged first by taking a single-frame snapshot. Dynein labeled with fluorophores (TMR or Alexa647) was imaged every 300 msec for 3 minutes. At the end, microtubules were imaged again by taking a snapshot to assess stage drift. Movies showing significant drift were not analyzed.

Kymographs were generated from motility movies, and dynein velocity and run length were calculated from kymographs using custom ImageJ macros as described.^74^ Only runs longer than 8 pixels were included in the analysis. Bright protein aggregates, which were defined as molecules 4x brighter than the background, were excluded. Velocity and run length were reported for individual dynein complexes. The processive landing rate was reported per microtubule by dividing the total number of processive events analyzed for that microtubule by the length of the microtubule, divided by the concentration of dynein used in that experimental condition, divided by the length of the movie in minutes. Five microtubules were analyzed per field of view for each condition. Data plotting and statistical analyses were performed in Prism9 (GraphPad).

### Binding assays with purified components

The binding affinity of Halo-CCSer2^650–850^ for Ndel1 was determined by coupling GST-Ndel1 to 30 μL of Glutathione Beads (Thermo Scientific) in 2 mL Protein Lo Bind Tubes (Eppendorf) using the following protocol. Beads were washed twice with 1 mL of GF150 without ATP supplemented with 10% glycerol and 0.1% NP-40. Ndel1 was diluted in this buffer to 0, 7.5, 15, 30, 60, 90, 120, and 300 nM in a final volume 25 μL, incubated with equilibrated beads, and gently shaken on a homemade device at room temperature for 1 h. 20 μL of supernatant were collected and analyzed via SDS-PAGE to confirm complete depletion of Ndel1. The Ndel1-conjugated beads were then washed once with 1 mL GF150 with 10% glycerol and 0.1% NP-40 and once with 1 mL of binding buffer (30 mM HEPES [pH 7.4], 2 mM magnesium acetate, 1 mM EGTA, 10% glycerol, 1 mM DTT, 1 mg/mL casein, 0.1% NP-40, and 1 mM ADP). The Halo-TMR labeled CCSer2^650–850^ was diluted to 5 nM in binding buffer and 25 μL was distributed to the Ndel1-conjugated beads. The mixture was gently agitated at room temperature for 45 minutes. After incubation, 20 μL of supernatant was collected and analyzed via SDS-PAGE to determine the depletion of Halo-CCSer2^650–850^. Depletion analysis was conducted through densitometry using ImageJ. Binding curves were fit in Prism10 (Graphpad) with a quadratic binding curve as described.^75^

The assay to assess Halo-CCSer2^650–850^ binding to dynein and Lis1 was performed similarly. Briefly, the beads were washed twice with 1 mL of GF150 supplemented with 0.1%NP40 and 25 uL Halo-CCSer2650-850 at 0, 300, 600, 900, or 1200 nM was added to the equilibrated beads and shaken for 1 hour. 20 μL of supernatant were collected and analyzed via SDS-PAGE. The beads were washed once with 1 mL GF150 with 10% glycerol and 0.1% NP-40 and once with 1 mL of binding buffer and 5 nM of either SNAP-549 or SNAP-647 dynein or SNAP-647 Lis1 in binding buffer was distributed to CCSer2-conjugated beads. The mixture was gently agitated at room temperature for 45 minutes. After incubation, 20 μL of supernatant was collected and analyzed via SDS-PAGE. Depletion analysis was conducted through densitometry using ImageJ.

Due to tag incompatibility, His-strep-sfGFP-CCSer2^650–850^ was used to assess CCSer2’s effect on dynein-Ndel1 or Lis1-Ndel1 interaction using a protocol similar to that described above. Briefly, 25 uL of Magne HaloTag Beads (Promega) were washed twice with 1 mL GF150 supplemented with 0.1%NP40. 5 nM Halo-Ndel1 in GF150 with 0.1%NP40 was added to the equilibrated beads and shaken for 1 hour for Lis1 experiments. For dynein experiments the final Ndel1 concentration was 30 nM. 20 μl of supernatant were collected and analyzed via SDS-PAGE. The beads were washed once with 1 mL GF150 with 10% glycerol and 0.1% NP-40 and once with 1 mL of binding buffer and 5 nM of either SNAP-549 or SNAP-647 dynein or SNAP-647 Lis1 in binding buffer with or without 90 nM strep-sfGFP-CCSer2^650–850^ was distributed to Ndel1-conjugated beads. The mixture was gently agitated at room temperature for 45 minutes. After incubation, 20 μL of supernatant was collected and analyzed via SDS-PAGE. Depletion analysis was conducted through densitometry using ImageJ.

### Data and software availability

All macros generated are available for download from the DeSantis-Lab GitHub page.

---

**Figure supplement 1:**
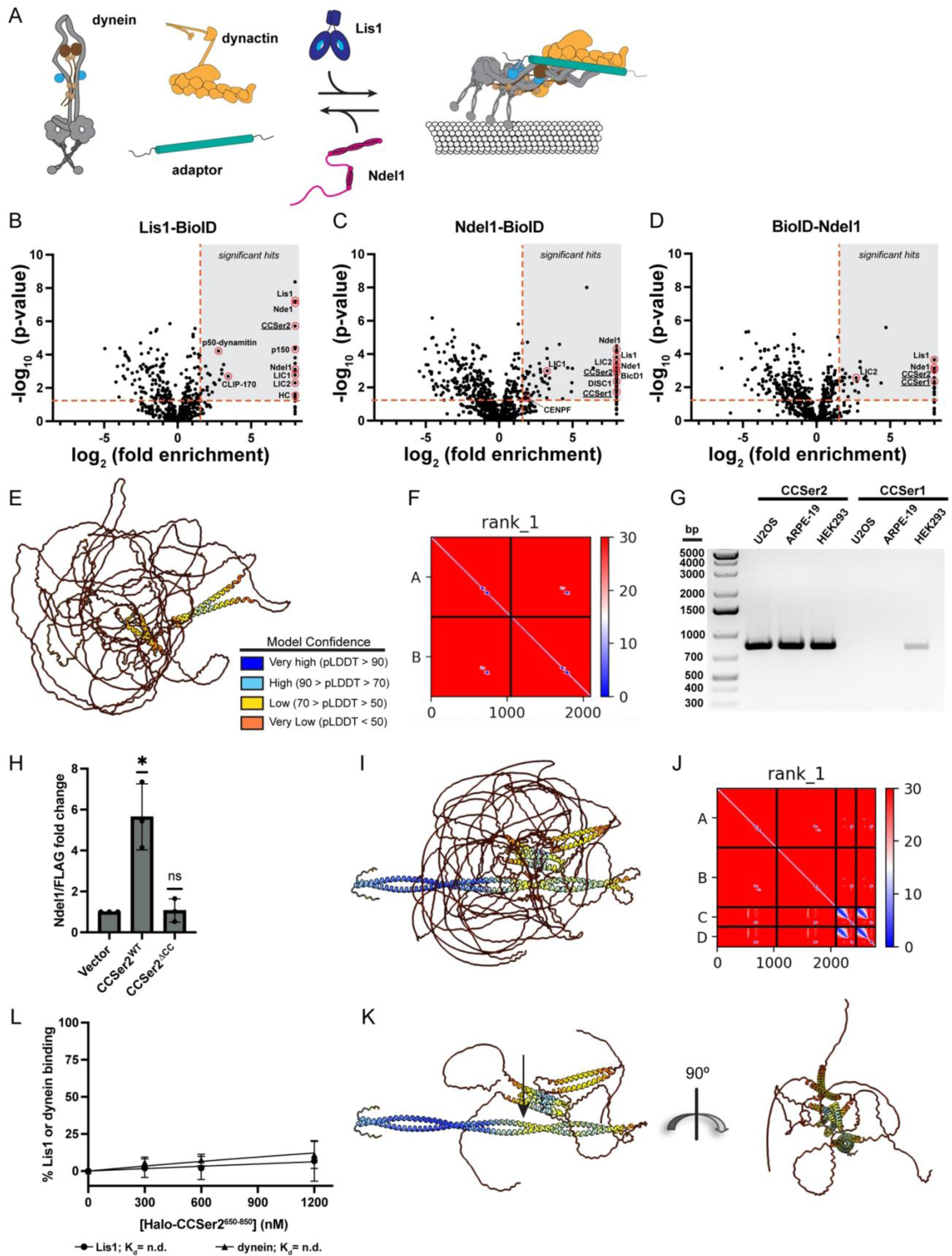
A) Schematic for dynein activation via binding dynactin and an adaptor. Lis1 promotes activation and Ndel1 inhibits. These proteins likely function in tandem in cells to support dynein activity though the mechanism of their tandem activity remains unclear. B-D) Volcano plots for Lis1-BioID (B), Ndel1-BioID (C), and BioID-Ndel1 (D). Enrichment over the BioID control versus significance between replicates are plotted. Significant hits used for interactome in (Figure 1A) are indicated by the grey box bounded by a p-value of 0.05 and a 3-fold enrichment over control (red dotted lines). Red circles denote proteins hits that are known interaction partners, as well as CCSer2 and CCSer1. Hits on the far right of the X-axis represent proteins that were not detected in the control. E) Alphafold prediction for two copies of CCSer2 colored by pLDDT as indicated. F) Predicted alignment error for the AlphaFold model shown in (E). G) PCR products of *ccser2* and *ccser1* amplified from cDNA libraries generated from the cell types indicated showing that CCSer2 is expressed to a higher extent and in more cell types than CCSer1. H) Quantification of Ndel1 co-immunoprecipitation with GFP vector, CCSer2^WT^, or CCSer2 ^ΔCC^. n = 3 biological replicates. The fold change was normalized to the amount of vector pulldown per experiment. Error bars shown are mean ± SD. Statistical analysis for each CCSer2^WT^ or CCSer2 ^ΔCC^ vs. Vector control was performed with one sample t and Wilcoxon test. I) Alphafold prediction of CCSer2 and Ndel1 colored by pLDDT score as indicated in Figure supplement 1E. J) Predicted alignment error for the AlphaFold model shown in (I). K) Model shown in (I) with disordered regions of CCSer2 removed for clarity. L) Binding curve between Lis1 (circle) or dynein (triangle) and CCSer2. n = 3 for both samples. Error bars mean ± SD.

**Figure supplement 2:**
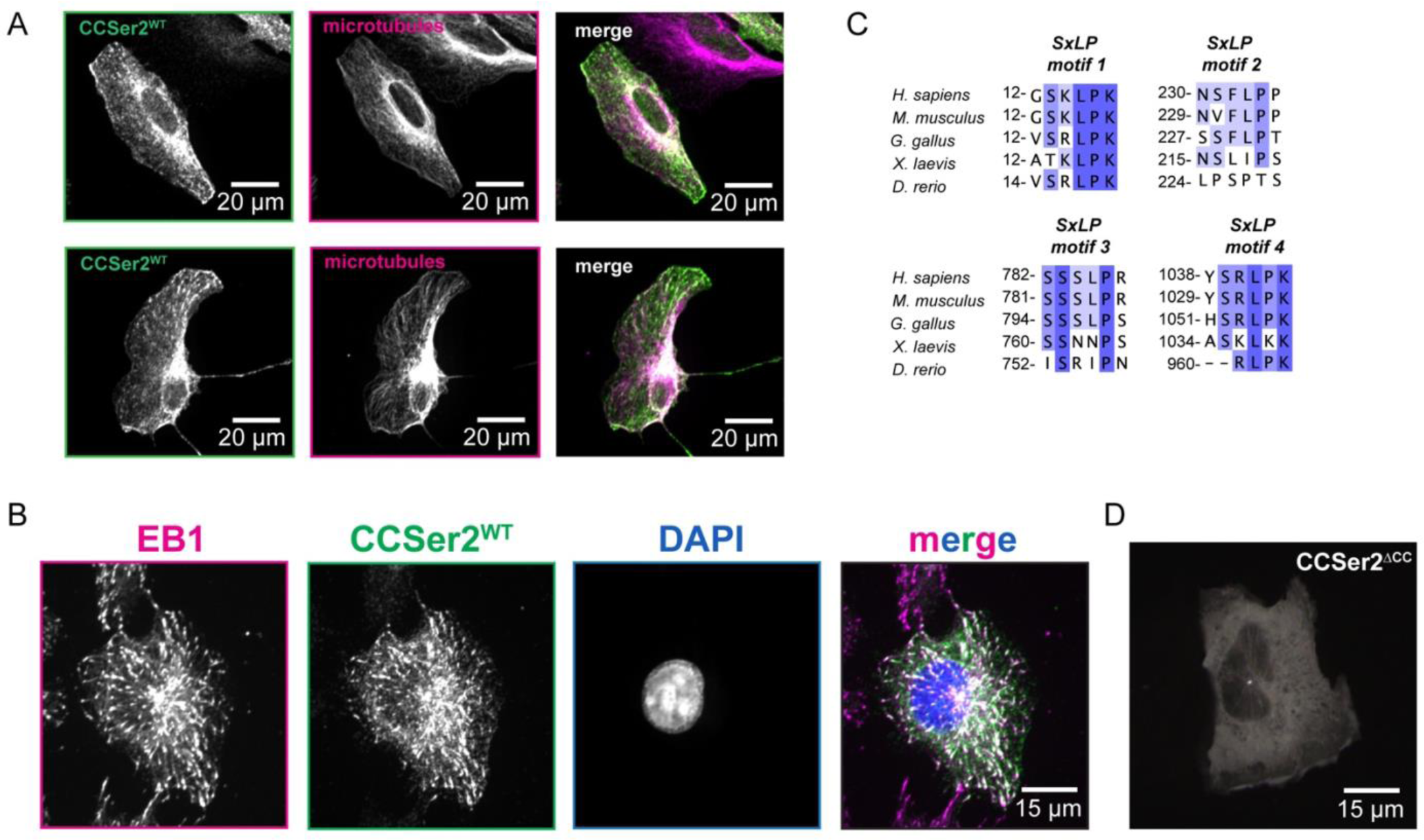
A) Additional representative images of U2OS cells stained for CCSer2^WT^ and microtubules as shown in Figure 1F. B) Fluorescence microscopy image fixed WT cells transfected with CCSer2^WT^ and stained with α-GFP to visualize CCSer2^WT^ (green), α-EB1 (magenta), and DAPI (blue). C) Alignment of CCSer2 S-x-L-P motifs 1 – 4. D) Temporal hyperstack colored as in Figure 3E for WT cells transfected with CCSer2 ^ΔCC^.

**Figure supplement 3:**
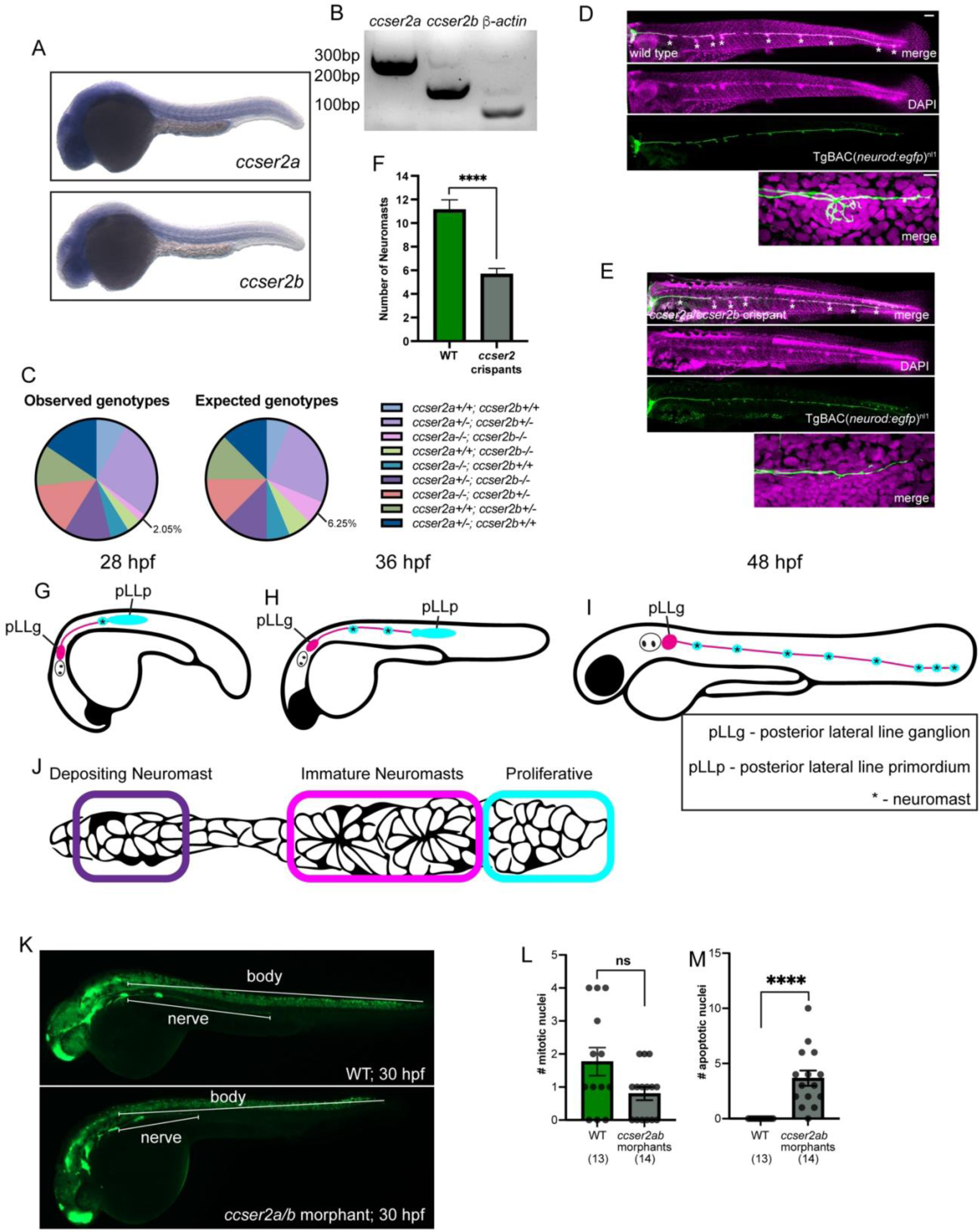
A) In situ hybridization showing *ccser2a* and *ccser2b* expression at 30 hpf in the zebrafish embryo. B) Maternal deposition of *ccser2a and ccser2b* mRNA compared to control B-actin mRNA, which is known to be maternally deposited at high levels. C) Observed and expected genotypes for *ccser2a;ccser2b* double crosses. D-E) DAPI (magenta) and neurons (green) of fixed wild type 4 dpf zebrafish larva (D) and *ccser2a/b* crispant (E). High magnification image of pLL nerve end shown below for each. Scale bar = 100 mm in whole trunk image; 10 mm in high magnification image below. F) Number of pLL neuromasts in wild type or *ccser2* crispants (Chi-square; p<0.005). G-J) Schematic showing pLL development (G-I) and organization of the pLL primordium (J). K) Measurement axis used for quantification of data in Figure 2E. L) Number of mitotic nuclei for wild type of *ccser2a/b* morphants (Chi-square; p=0.2514). M) Number of cells undergoing apoptosis for wild type and *ccser2a/b* crispants (Chi-square; p<0.005).

**Figure supplement 4:**
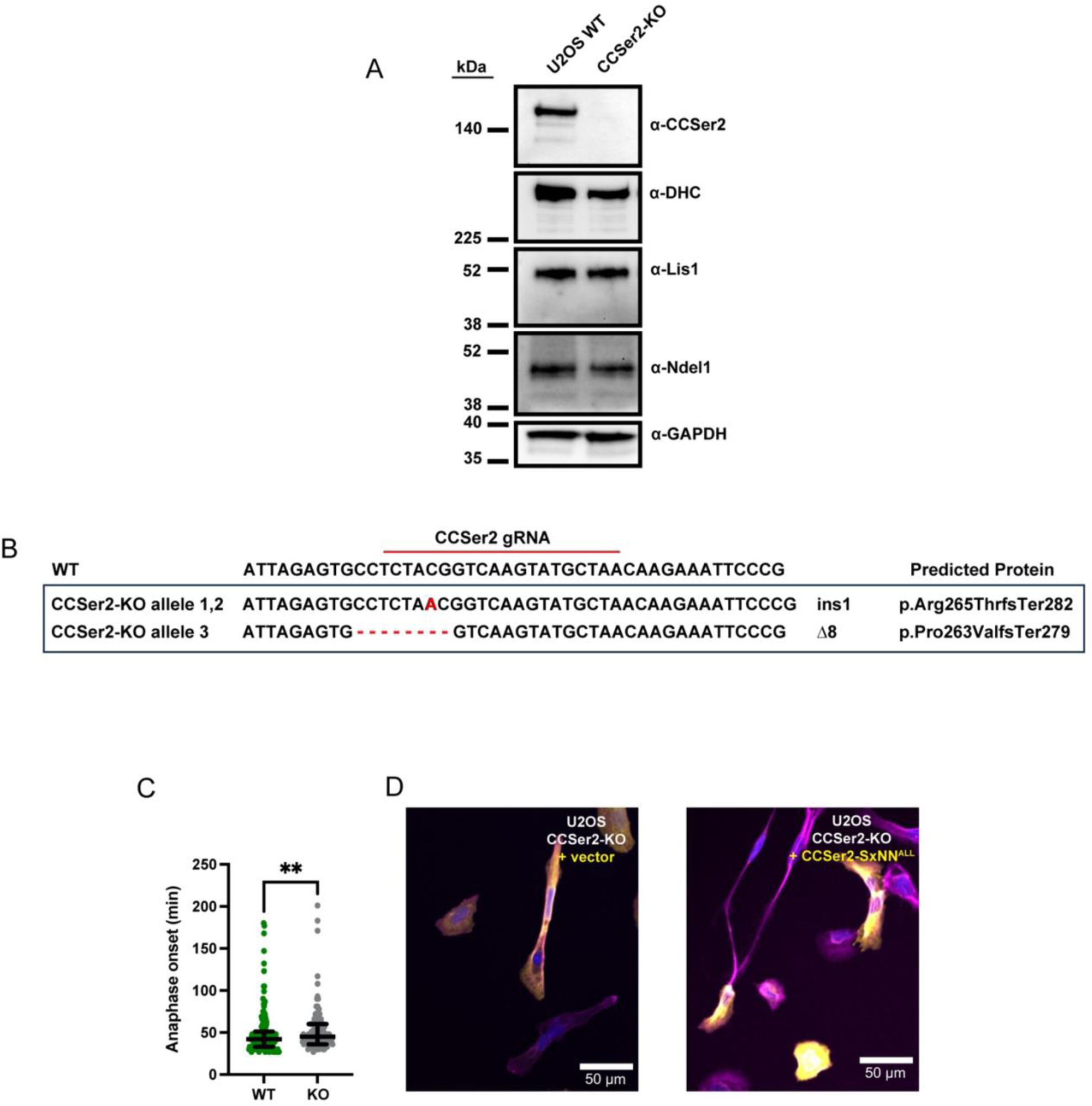
A) Western blot of lysate from WT cells and CCSer2-KO cell lines. Membranes were probed with α-CCSer2, α-DHC, α-Lis1, α-Ndel1, and α-GAPDH as a loading control. B) Results of indel analysis of the CCSer2-KO. Two frameshift (fs) mutations identified in the CCSer2 gene result in early stop codons and truncation of the protein (Ter282 and Ter279). The single base pair insertion mutation occurred 80% of the alleles sequenced (24/30 alleles) and the 8 base pair deletion mutation occurred in 20% of the alleles sequenced (6/30 alleles). No WT alleles were found in the CCSer2-KO. C) The time to anaphase of dividing WT and CCSer2-KO cells is quantified. n = 170 and 102 mitotic cells analyzed for WT and CCSer2-KOs, respectively. Error bars are median ± interquartile range. Statistical analysis performed with a Mann-Whitney test. D) Fluorescence microscopy images of fixed CCSer2-KOs transfected with GFP vector or CCSer2-SxNN^ALL^. Cells were stained with phalloidin to visualize actin (pink), α-GFP to visualize vector and CCSer2-SXNN^ALL^ (yellow), and DAPI to visualize nuclei (blue). The average circularity of the samples shown in (D) is quantified in Figure 2H.

**Figure supplement 5:**
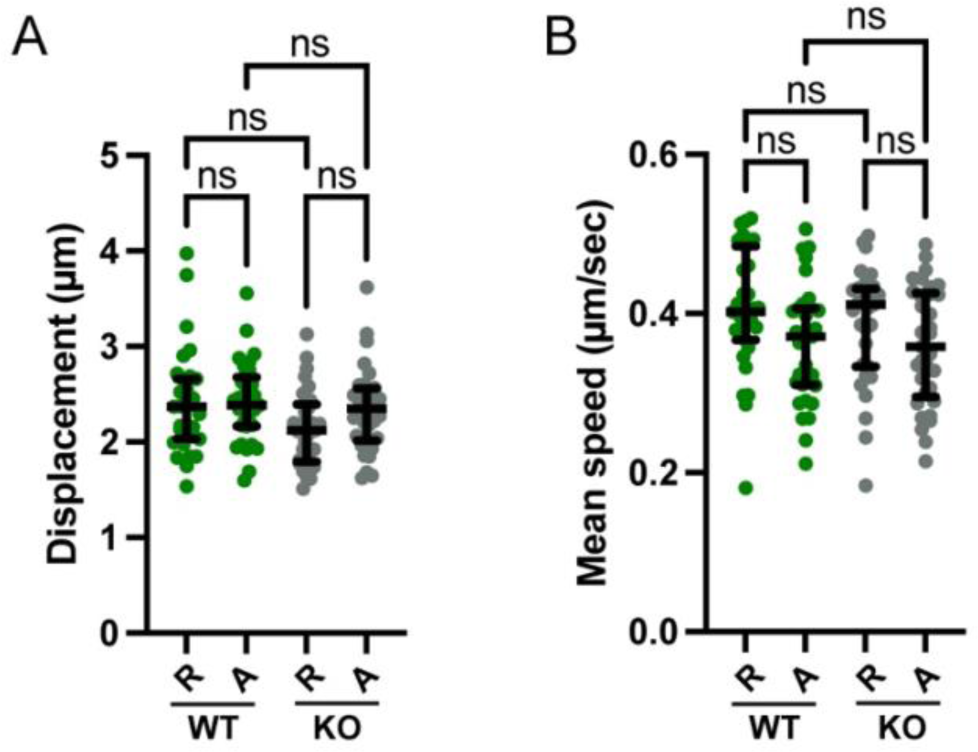
A) The displacement of the averaged anterograde and retrograde m-Cherry integrin tracks per cell of WT and CCSer2-KOs. n = 31 and 33 cells analyzed for WT and CCSer2-KO cells, respectively, across three biological replicates. B) The mean speed of the averaged anterograde and retrograde integrin tracks per cell of WT and CCSer2-KOs are not significantly different. n = 31 and 33 cells analyzed for WT and CCSer2-KO cells, respectively, across three biological replicates. A, B) Error bars are median ± interquartile range. Statistical analysis performed with a Kruskal-Wallis test with Dunn’s multiple comparisons test.

**Figure supplement 6:**
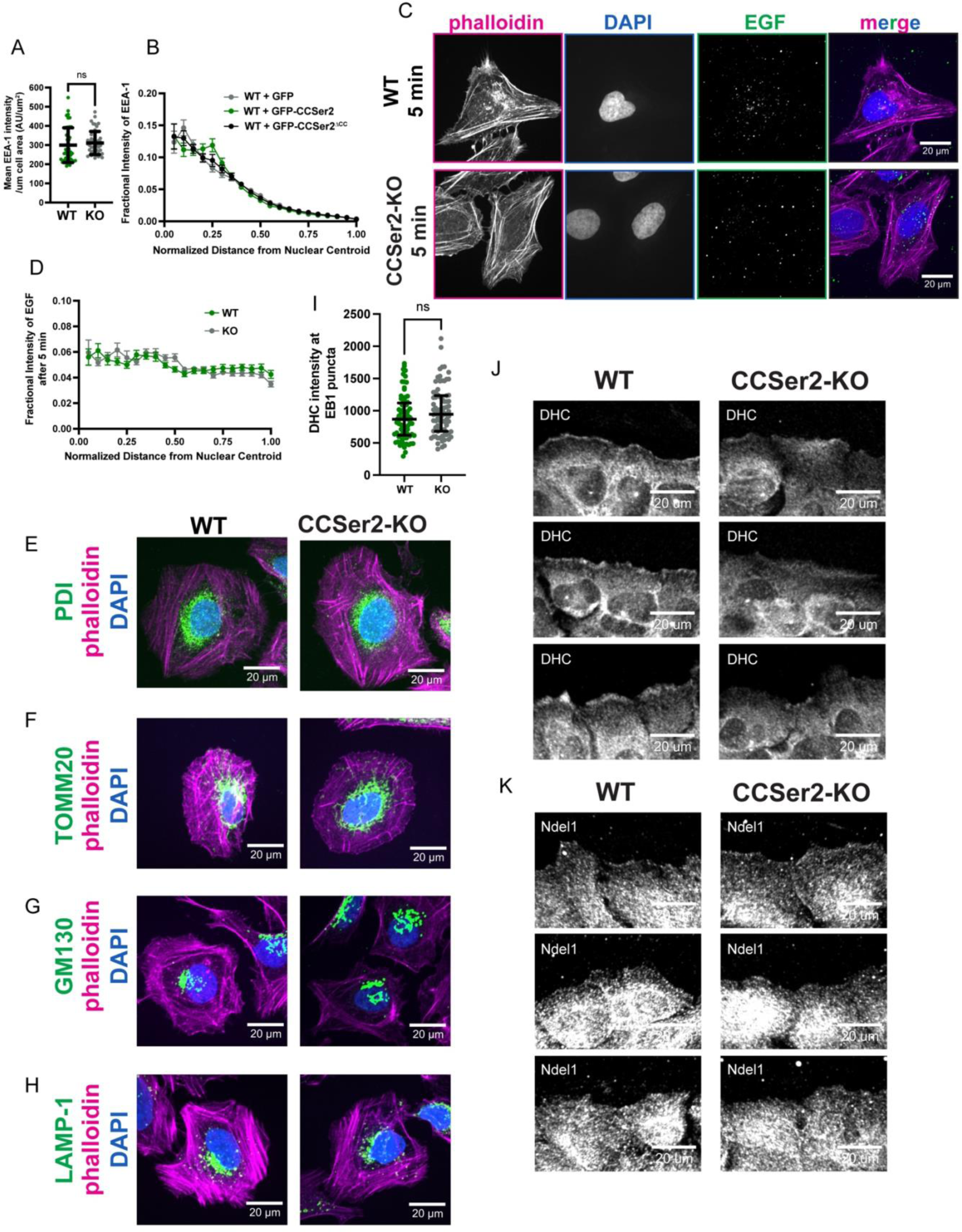
A) Mean intensity of early endosomes per cell area. n = 46 cells analyzed per sample across three biological replicates. Error bars are median ± interquartile range. Statistical analysis performed with a Mann-Whitney test. B) The quantification of the fractional early endosome intensity from the centroid of the nucleus to the cell periphery in WT cells transfected with vector, CCSer2^WT^, or CCSer2^ΔCC^ shows no significant differences in the relative distribution of early endosomes. n = 44-46 cells analyzed per sample across three biological replicates. C) Fluorescence microscopy images of fixed WT and CCSer2-KO cells treated with EGF-555, shown in green, for 5 minutes and excess washed out. Cells were stained with phalloidin to visualize actin (magenta) and DAPI to visualize nuclei (blue). After 5 minutes of EGF-555 treatment, neither WT nor CCSer2-KO cells have yet to cluster EGF+ endosomes. C) The quantification of the fractional EGF-555 intensity from the centroid of the nucleus to the cell periphery in WT and CCSer2-KOs five minutes after treatment shows no preferential enrichment of EGF+ endosomes in any region of the cell. n = 89 and 85 cells analyzed for WT and CCSer2-KOs, respectively, across three biological replicates. B, D) Error bars shown are mean ± SEM. Statistical analysis performed with multiple, Mann-Whitney tests and an FDR of 1%. E-H) Fluorescence microscopy images of PFA-fixed WT and CCSer2-KO cells stained with phalloidin to visualize actin (magenta), DAPI to visualize nuclei (blue), and antibodies against the following organelles and shown in green: E) ER (α-PDI), E) mitochondria (α-TOMM20), F) Golgi apparatus (α-GM130), and G) lysosomes (α-LAMP-1). Quantification is shown in Figure 4 H-K. I) Intensity of dynein (stained with α-dynein heavy chain) at EB1 puncta in WT and CCSer2-KO cells. Error bars are median with interquartile range. Significance determined with a Mann-Whitney test. J, K) Additional representative images of WT and CCSer2 KO cells fixed 30 minutes post wounding and stained with α-dynein heavy chain (α-DHC) (J) or α-Ndel1 (K) as shown in Figure 4L, N.

**Figure supplement 7:**
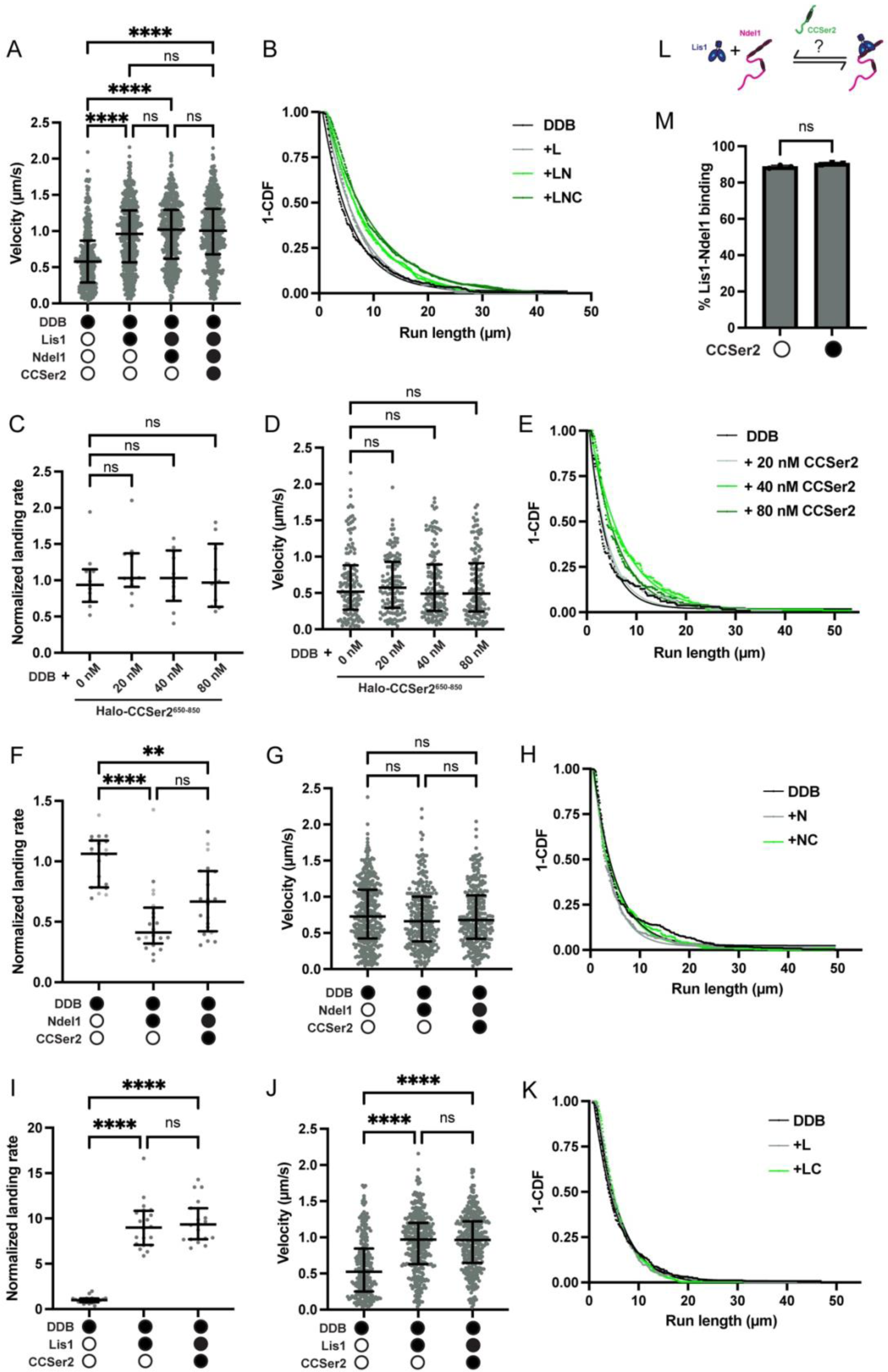
A) Velocity of dynein-dynactin-BicD2 (DDB) complexes in the absence (white circles) or presence (black circles) of 50 nM Lis1, 10 nM Ndel1, and/or 80 nM CCSer2^650–850^. B) 1-Cumulative distribution frequency of run length for dynein-dynactin-BicD2 complexes (DDB) with combinations of Lis1, Ndel1, and CCSer2^650–850^. C) Number of processive events per µm of microtubule per nM of dynein for dynein-dynactin-BicD2 (DDB) complexes in the presence of increasing concentrations of Halo-CCSer2^650–850^ as indicated. D) Velocity of dynein-dynactin-BicD2 (DDB) complexes in the presence of increasing concentrations of CCSer2^650–850^ as indicated. E) 1-Cumulative distribution frequency of run length for dynein-dynactin-BicD2 complexes (DDB) with increasing concentrations of CCSer2^650–850^ as indicated. F) Number of processive events per µm of microtubule per nM of dynein for dynein-dynactin-BicD2 (DDB) complexes in the absence (white circles) or presence (black circles) of 10 nM Ndel1 and 80nM Halo-CCSer2^650–850^. G) Velocity of dynein-dynactin-BicD2 (DDB) in the in the absence (white circles) or presence (black circles) of 10 nM Ndel1 or 80 nM Halo-CCSer2^650–850^. H) 1 - Cumulative distribution frequency of run length for dynein-dynactin-BicD2 complexes (DDB) alone or with 10 nM Ndel1 alone and 80 nM Halo-CCSer2^650–850^. I) Number of processive events per µm of microtubule per nM of dynein for dynein-dynactin-BicD2 (DDB) complexes in the absence (white circles) or presence (black circles) of 50 nM Lis1 or 80nM CCSer2^650–850^. J) Velocity of dynein-dynactin-BicD2 (DDB) in the absence (white circles) or presence (black circles) of 5 nM Lis1 and 80nM CCSer2^650–850^. K) 1 - Cumulative distribution frequency of run length for dynein-dynactin-BicD2 complexes (DDB) alone or with 50 nM Lis1 and with 80 nM CCSer2^650–850^. A, D, G, J) Error bars are median ± interquartile range. Statistical analysis performed with a Kruskal-Wallis test with Dunn’s multiple comparisons test. C, F, I) Data are normalized to the dynein-dynactin-BicD2 sample for each replicate. n = 25 microtubules analyzed for each condition. Statistical analysis performed with a Brown-Forsythe and Welch ANOVA with Dunnet’s T3 multiple comparison test. L) Schematic of assay to determine if CCSer2 affects Lis1-Ndel1 binding. M) Percent of Lis1 bound to Ndel1-conjugated beads in the absence (white circle) or presence (black circle) of Halo-CCSer2^650–850^. n = 3. Statistical analysis performed with a Mann Whitney test.

## Legends for supplemental movies

**Supplemental movie-1.** Related to Figure 1. Representative movie of U2OS WT cell expressing CCSer2^WT^. Cells were imaged 48 hours post transfection with a 60X objective at 1 frame per second.

**Supplemental movie-2.** Related to Figure 1. Movie of U2OS WT cell expressing CCSer2-SxNN^ALL^. Cells were imaged 48 hours post transfection with a 60X objective at 1 frame per second. Many cells CCSer2-SxNN^ALL^ had fewer comets or lower expression levels.

**Supplemental movies-3, 4.** Related to Figure 2. Representative movies of U2OS WT cells plated on fibronectin and imaged at 20X with a frame rate of 1 frame every 3 minutes. Arrows designate lamellipodia of individual migrating cells.

**Supplemental movies-5,6.** Related to Figure 2. Representative movies of U2OS CCSer2-KO cells plated on fibronectin and imaged at 20X with a frame rate of 1 frame every 3 minutes.

**Supplemental movies-7,8.** Related to Figure 42. Representative movies of U2OS WT or CCSer2-KO cells, respectively, plated on fibronectin around 2-well silicone inserts. Nuclei labeled with SiR-DNA dye. 10X imaging began approx. an hour after removal of the silicone insert with a frame rate of 1 frame every 3 minutes.

## References

1. Bodakuntla, S., Jijumon, A. S., Villablanca, C., Gonzalez-Billault, C. & Janke, C. Microtubule-Associated Proteins: Structuring the Cytoskeleton. Trends in Cell Biology 29, 804–819 (2019).

2. Hirokawa, N., Noda, Y., Tanaka, Y. & Niwa, S. Kinesin superfamily motor proteins and intracellular transport. Nat Rev Mol Cell Biol 10, 682–696 (2009).

3. Cianfrocco, M. A., DeSantis, M. E., Leschziner, A. E. & Reck-Peterson, S. L. Mechanism and Regulation of Cytoplasmic Dynein. Annu. Rev. Cell Dev. Biol. 31, 83–108 (2015).

4. Miki, H., Okada, Y. & Hirokawa, N. Analysis of the kinesin superfamily: insights into structure and function. Trends in Cell Biology 15, 467–476 (2005).

5. Reck-Peterson, S. L., Redwine, W. B., Vale, R. D. & Carter, A. P. The cytoplasmic dynein transport machinery and its many cargoes. Nature Reviews Molecular Cell Biology 19, 382–398 (2018).

6. Schlager, M. A., Hoang, H. T., Urnavicius, L., Bullock, S. L. & Carter, A. P. In vitro reconstitution of a highly processive recombinant human dynein complex. EMBO J 33, 1855–1868 (2014).

7. McKenney, R. J., Huynh, W., Tanenbaum, M. E., Bhabha, G. & Vale, R. D. Activation of cytoplasmic dynein motility by dynactin-cargo adapter complexes. Science 345, 337–341 (2014).

8. Garrott, S. R., Gillies, J. P. & DeSantis, M. E. Nde1 and Ndel1: Outstanding Mysteries in Dynein-Mediated Transport. Front Cell Dev Biol 10, 871935 (2022).

9. Markus, S. M., Marzo, M. G. & McKenney, R. J. New insights into the mechanism of dynein motor regulation by lissencephaly-1. eLife 9, e59737 (2020).

10. Coquelle, F. M. et al. LIS1, CLIP-170’s Key to the Dynein/Dynactin Pathway. Molecular and Cellular Biology 22, 3089–3102 (2002).

11. Splinter, D. et al. BICD2, dynactin, and LIS1 cooperate in regulating dynein recruitment to cellular structures. MBoC 23, 4226–4241 (2012).

12. Sitaram, P., Anderson, M. A., Jodoin, J. N., Lee, E. & Lee, L. A. Regulation of dynein localization and centrosome positioning by Lis-1 and asunder during Drosophila spermatogenesis. Development 139, 2945–2954 (2012).

13. Stehman, S. A., Chen, Y., McKenney, R. J. & Vallee, R. B. NudE and NudEL are required for mitotic progression and are involved in dynein recruitment to kinetochores. J Cell Biol 178, 583–594 (2007).

14. Liang, Y. et al. Nudel Modulates Kinetochore Association and Function of Cytoplasmic Dynein in M Phase. Mol Biol Cell 18, 2656–2666 (2007).

15. Wynne, C. L. & Vallee, R. B. Cdk1 phosphorylation of the dynein adapter Nde1 controls cargo binding from G2 to anaphase. J Cell Biol 217, 3019–3029 (2018).

16. Raaijmakers, J. A., Tanenbaum, M. E. & Medema, R. H. Systematic dissection of dynein regulators in mitosis. J Cell Biol 201, 201–215 (2013).

17. Htet, Z. M. et al. LIS1 promotes the formation of activated cytoplasmic dynein-1 complexes. Nature Cell Biology 22, 518–525 (2020).

18. Elshenawy, M. M. et al. Lis1 activates dynein motility by modulating its pairing with dynactin. Nat Cell Biol 22, 570–578 (2020).

19. Gutierrez, P. A., Ackermann, B. E., Vershinin, M. & McKenney, R. J. Differential effects of the dynein-regulatory factor Lissencephaly-1 on processive dynein-dynactin motility. J Biol Chem 292, 12245–12255 (2017).

20. Okada, K. et al. Conserved Roles for the Dynein Intermediate Chain and Ndel1 in Assembly and Activation of Dynein. 2023.01.13.523097 Preprint at 10.1101/2023.01.13.523097 (2023).

21. Garrott, S. R. et al. Ndel1 disfavors dynein–dynactin–adaptor complex formation in two distinct ways. Journal of Biological Chemistry 299, (2023).

22. McKenney, R. J., Weil, S. J., Scherer, J. & Vallee, R. B. Mutually Exclusive Cytoplasmic Dynein Regulation by NudE-Lis1 and Dynactin. J. Biol. Chem. 286, 39615–39622 (2011).

23. Zhao, Y., Oten, S. & Yildiz, A. Nde1 Promotes Lis1-Mediated Activation of Dynein. 2023.05.26.542537 Preprint at 10.1101/2023.05.26.542537 (2023).

24. Żyłkiewicz, E., et al. The N-terminal coiled-coil of Ndel1 is a regulated scaffold that recruits LIS1 to dynein. J Cell Biol 192, 433–445 (2011).

25. Roux, K. J., Kim, D. I. & Burke, B. BioID: A Screen for Protein-Protein Interactions: BioID Screen for Protein Interactions. in Current Protocols in Protein Science (eds. Coligan, J. E., Dunn, B. M., Speicher, D. W. & Wingfield, P. T.) 19.23.1–19.23.14 (John Wiley & Sons, Inc., Hoboken, NJ, USA, 2013). doi:10.1002/0471140864.ps1923s74.

26. Redwine, W. B. et al. The human cytoplasmic dynein interactome reveals novel activators of motility. Elife 6, (2017).

27. Bradshaw, N. J., Hennah, W. & Soares, D. C. NDE1 and NDEL1: twin neurodevelopmental proteins with similar ‘nature’ but different ‘nurture’. BioMolecular Concepts 4, (2013).

28. Soares, D. C. et al. The Mitosis and Neurodevelopment Proteins NDE1 and NDEL1 Form Dimers, Tetramers, and Polymers with a Folded Back Structure in Solution *. Journal of Biological Chemistry 287, 32381–32393 (2012).

29. Grantham, J. The Molecular Chaperone CCT/TRiC: An Essential Component of Proteostasis and a Potential Modulator of Protein Aggregation. Frontiers in Genetics 11, (2020).

30. Wang, M., Herrmann, C. J., Simonovic, M., Szklarczyk, D. & von Mering, C. Version 4.0 of PaxDb: Protein abundance data, integrated across model organisms, tissues, and cell-lines. Proteomics 15, 3163–3168 (2015).

31. Jumper, J. et al. Highly accurate protein structure prediction with AlphaFold. Nature 596, 583–589 (2021).

32. Mirdita, M. et al. ColabFold: making protein folding accessible to all. Nat Methods 19, 679–682 (2022).

33. Mun, D. J. et al. Gcap14 is a microtubule plus-end-tracking protein coordinating microtubule–actin crosstalk during neurodevelopment. Proceedings of the National Academy of Sciences 120, e2214507120 (2023).

34. Honnappa, S. et al. An EB1-binding motif acts as a microtubule tip localization signal. Cell 138, 366–376 (2009).

35. Jiang, K. et al. A Proteome-wide screen for mammalian SxIP motif-containing microtubule plus-end tracking proteins. Curr. Biol. 22, 1800–1807 (2012).

36. El-Brolosy, M. A. et al. Genetic compensation triggered by mutant mRNA degradation. Nature 568, 193– 197 (2019).

37. Shah, A. N., Davey, C. F., Whitebirch, A. C., Miller, A. C. & Moens, C. B. Rapid reverse genetic screening using CRISPR in zebrafish. Nat Methods 12, 535–540 (2015).

38. Luxton, G. G. & Gundersen, G. G. Orientation and function of the nuclear–centrosomal axis during cell migration. Current Opinion in Cell Biology 23, 579–588 (2011).

39. Schmoranzer, J. et al. Par3 and Dynein Associate to Regulate Local Microtubule Dynamics and Centrosome Orientation during Migration. Current Biology 19, 1065–1074 (2009).

40. Etienne-Manneville, S. & Hall, A. Integrin-mediated activation of Cdc42 controls cell polarity in migrating astrocytes through PKCzeta. Cell 106, 489–498 (2001).

41. Palazzo, A. F. et al. Cdc42, dynein, and dynactin regulate MTOC reorientation independent of Rho-regulated microtubule stabilization. Current Biology 11, 1536–1541 (2001).

42. Levy, J. R. & Holzbaur, E. L. F. Dynein drives nuclear rotation during forward progression of motile fibroblasts. Journal of Cell Science 121, 3187–3195 (2008).

43. Shafaq-Zadah, M. et al. Persistent cell migration and adhesion rely on retrograde transport of β 1 integrin. Nature Cell Biology 18, 54–64 (2016).

44. Baumbach, J. et al. Lissencephaly-1 is a context-dependent regulator of the human dynein complex. eLife 6, e21768 (2017).

45. Sasaki, S. et al. A LIS1/NUDEL/Cytoplasmic Dynein Heavy Chain Complex in the Developing and Adult Nervous System. Neuron 28, 681–696 (2000).

46. Taberner, N. & Dogterom, M. Motor-mediated clustering at microtubule plus ends facilitates protein transfer to a bio-mimetic cortex. 736728 Preprint at 10.1101/736728 (2019).

47. Siegrist, S. E. & Doe, C. Q. Microtubule-induced cortical cell polarity. Genes Dev. 21, 483–496 (2007).

48. Moon, H. M. et al. LIS1 controls mitosis and mitotic spindle organization via the LIS1–NDEL1–dynein complex. Human Molecular Genetics 23, 449–466 (2014).

49. Yamada, M. et al. LIS1 and NDEL1 coordinate the plus-end-directed transport of cytoplasmic dynein. EMBO J 27, 2471–2483 (2008).

50. Lee, W.-L., Oberle, J. R. & Cooper, J. A. The role of the lissencephaly protein Pac1 during nuclear migration in budding yeast. J Cell Biol 160, 355–364 (2003).

51. Moughamian, A. J., Osborn, G. E., Lazarus, J. E., Maday, S. & Holzbaur, E. L. F. Ordered Recruitment of Dynactin to the Microtubule Plus-End is Required for Efficient Initiation of Retrograde Axonal Transport. Journal of Neuroscience 33, 13190–13203 (2013).

52. Fellows, A. D., Bruntraeger, M., Burgold, T., Bassett, A. R. & Carter, A. P. Dynein and dynactin move long-range but are delivered separately to the axon tip. 2023.07.03.547521 Preprint at 10.1101/2023.07.03.547521 (2023).

53. Ogawa, F. et al. NDE1 and GSK3β Associate with TRAK1 and Regulate Axonal Mitochondrial Motility: Identification of Cyclic AMP as a Novel Modulator of Axonal Mitochondrial Trafficking. ACS Chem. Neurosci. 7, 553–564 (2016).

54. Kuijpers, M. et al. Dynein Regulator NDEL1 Controls Polarized Cargo Transport at the Axon Initial Segment. Neuron 89, 461–471 (2016).

55. Ye, F. et al. DISC1 regulates neurogenesis via modulating kinetochore attachment of Ndel1/Nde1 during mitosis. Neuron 96, 1041–1054.e5 (2017).

56. Ye, J. et al. Mechanistic insights into the interactions of dynein regulator Ndel1 with neuronal ankyrins and implications in polarity maintenance. Proc Natl Acad Sci U S A 117, 1207–1215 (2020).

57. Pouthas, F. et al. In migrating cells, the Golgi complex and the position of the centrosome depend on geometrical constraints of the substratum. Journal of Cell Science 121, 2406–2414 (2008).

58. Shafaq-Zadah, M., Dransart, E. & Johannes, L. Clathrin-independent endocytosis, retrograde trafficking, and cell polarity. Curr Opin Cell Biol 65, 112–121 (2020).

59. Bielska, E. et al. Hook is an adapter that coordinates kinesin-3 and dynein cargo attachment on early endosomes. J Cell Biol 204, 989–1007 (2014).

60. Kendrick, A. A. et al. Hook3 is a scaffold for the opposite-polarity microtubule-based motors cytoplasmic dynein-1 and KIF1C. J Cell Biol 218, 2982–3001 (2019).

61. Gibson, D. G. et al. Enzymatic assembly of DNA molecules up to several hundred kilobases. Nat Methods 6, 343–345 (2009).

62. Stemmer, M., Thumberger, T., Keyer, M. del S., Wittbrodt, J. & Mateo, J. L. CCTop: An Intuitive, Flexible and Reliable CRISPR/Cas9 Target Prediction Tool. PLOS ONE 10, e0124633 (2015).

63. Ran, F. A. et al. Genome engineering using the CRISPR-Cas9 system. Nat Protoc 8, 2281–2308 (2013).

64. Yamaji, T. & Hanada, K. Establishment of HeLa Cell Mutants Deficient in Sphingolipid-Related Genes Using TALENs. PLOS ONE 9, e88124 (2014).

65. Ershov, D. et al. TrackMate 7: integrating state-of-the-art segmentation algorithms into tracking pipelines. Nat Methods 19, 829–832 (2022).

66. Liang, C.-C., Park, A. Y. & Guan, J.-L. In vitro scratch assay: a convenient and inexpensive method for analysis of cell migration in vitro. Nat Protoc 2, 329–333 (2007).

67. ZFIN Publication: Westerfield, 2007. https://zfin.org/ZDB-PUB-101222-53.

68. Kimmel, C. B., Ballard, W. W., Kimmel, S. R., Ullmann, B. & Schilling, T. F. Stages of embryonic development of the zebrafish. Dev Dyn 203, 253–310 (1995).

69. Thisse, C. & Thisse, B. High-resolution in situ hybridization to whole-mount zebrafish embryos. Nat Protoc 3, 59–69 (2008).

70. Logel, J., Dill, D. & Leonard, S. Synthesis of cRNA probes from PCR-generated DNA. Biotechniques 13, 604–610 (1992).

71. Agrawal, R. et al. The KASH5 protein involved in meiotic chromosomal movements is a novel dynein activating adaptor. eLife 11, e78201 (2022).

72. Zhang, K. et al. Cryo-EM Reveals How Human Cytoplasmic Dynein Is Auto-inhibited and Activated. Cell 169, 1303–1314.e18 (2017).

73. Huang, J., Roberts, A. J., Leschziner, A. E. & Reck-Peterson, S. L. Lis1 Acts as a “Clutch” between the ATPase and Microtubule-Binding Domains of the Dynein Motor. Cell 150, 975–986 (2012).

74. Roberts, A. J., Goodman, B. S. & Reck-Peterson, S. L. Reconstitution of dynein transport to the microtubule plus end by kinesin. eLife 3, e02641 (2014).

75. Pollard, T. D. A Guide to Simple and Informative Binding Assays. MBoC 21, 4061–4067 (2010).

